# Towards the accurate modelling of antibody-antigen complexes from sequence using machine learning and information-driven docking

**DOI:** 10.1101/2023.11.17.567543

**Authors:** Marco Giulini, Constantin Schneider, Daniel Cutting, Nikita Desai, Charlotte M. Deane, Alexandre M.J.J. Bonvin

## Abstract

Antibody-antigen complex modelling is an important step in computational workflows for therapeutic antibody design. While experimentally determined structures of both antibody and the cognate antigen are often not available, recent advances in machine learning-driven protein modelling have enabled accurate prediction of both antibody and antigen structures. Here, we analyse the ability of protein-protein docking tools to use machine learning generated input structures for information-driven docking. We find that HADDOCK can generate accurate models of antibody-antigen complexes using an ensemble of antibody structures generated by machine learning tools and AlphaFold2 predicted antigen structures. Targeted docking using knowledge of the complementary determining regions on the antibody and some information about the targeted epitope allows the generation of high quality models of the complex with reduced sampling, resulting in a computationally cheap protocol that outperforms the ZDOCK baseline. The data set used to benchmark the docking protocols in this study is available at github.com/haddocking/ai-antibodies. The docking models will be deposited at data.sbgrid.org/labs/32/ upon acceptance.

## I. INTRODUCTION

Antibodies are Y-shaped proteins produced by B-cells that bind with high selectivity and affinity to invading antigens recognized as potentially dangerous by the immune system, making them useful candidates for therapeutics development: as of June 2022, 162 antibody therapeutics have been approved globally [1]. Their highly desirable binding properties are mainly due to the process of somatic hypermutation, in which the complementarity determining regions (CDRs) of the antibody are optimized in order to specifically bind the epitope, the set of amino acids on the antigen molecule engaged by the antibody. The CDRs of an antibody are six hypervariable loops, distributed over the variable regions of the light and heavy chains. The third hypervariable loop on the heavy chain (CDR H3) usually corresponds to the most important region for antigen binding [2], as well as the most difficult region to be modelled by computational approaches [3–5], due to its high variability in both sequence composition and length. As of 30 Oct 2023, 7853 antibody structures have been collated in the Structural Antibody Database (SAbDab) [6], the majority of which, 7495, are found in complex with the cognate antigen.

Following the recent achievements in computer-aided structure prediction [7–9], several studies have shown that it is possible to accurately model antibody structures from sequence information using machine learning (ML)-based approaches [5, 10].

While ML-based antibody modelling tools are already reaching high accuracy [11], the accurate prediction of antibody-antigen complex structures from sequence is a much harder problem, which still represents a challenge for state-of-the-art methods such as AlphaFold2-Multimer [12, 13]. As an example, the three antibody-antigen complexes present in the recent CASP15-CAPRI54 challenges [14, 15] proved to be among the most challenging targets [16]. In this context, physics-based, information-driven docking algorithms are useful to generate reasonable poses for the desired complex and thus form part of the backbone of many current antibody design workflows [17–19].

Here, we demonstrate that protocols combining ML-driven antibody and antigen modelling with information-driven docking can produce useful docking poses without access to experimentally determined antibody or antigen structures. We first demonstrate that AlphaFold2-Multimer [12] has limited accuracy when used to model antibody-antigen complexes. We then compare four different ML-based antibody modelling tools, ABodyBuilder2 [5], ABlooper [20], AlphaFold2-Multimer, and IgFold [10], to assess their ability to generate antibody models that are accurate enough to be suitable to be fed into our information-driven docking software, HADDOCK [21], with the aim of generating accurate models of the antibody-antigen complex. In order to mimic realistic scenarios, we consider not only the experimentally determined (“bound”) structure of the antigen, but also AlphaFold2 [8, 12] models of the target antigen from sequence.

We show that, even limiting the sampling of solutions to a few tens of models, HADDOCK is capable of yielding acceptable models of the antibody-antigen complex. By comparing different docking protocols, we identify model diversity in the input antibody structures as the key for higher docking success. We highlight a protocol combining an ensemble of antibody structures generated by ABodyBuilder2 and antigen structures predicted by AlphaFold2, which, using information-driven HADDOCK, models antibody-antigen complexes with acceptable accuracy without relying on experimentally determined structures of either interaction partner and is computationally inexpensive.

## II. METHODS

### A. Data set construction

We constructed a benchmark data set to evaluate the docking performance of different protocols on both experimentally determined and computationally modelled structures of antibodies and cognate antigens. Crucially, previously used benchmark data sets (such as the data set used by Ambrosetti *et al*. [4]) are not well suited to the assessment of ML-driven antibody modeling and docking workflows, since data included in these data sets will have been included in training sets for current state-of-the-art ML methods. We therefore used SAbDab [6] to generate a data set of antibody-antigen complex structures which fulfilled the following criteria:

1. Each structure in the data set had to be deposited after the training of the ML-driven structural modeling tools we used to predict antibody and antigen structures. We used July 2021 as the cutoff date for additions to the data set.
2. Antibodies in the data set should not share their concatenated CDR sequences (defined using IMGT definition [22]) with any antibody deposited before the creation of the benchmark data set nor any other antibody in the benchmark data set.
3. The experimentally determined structure should have a resolution *≤* 3 Å.
4. Only complexes with protein antigens (excluding peptides under 30 amino acids length) were considered.
5. Only complete Fabs were added to the data set, nanobodies were excluded.

The resulting data set contains 71 antibody-antigen complexes (see Supplementary Table 1).

### B. Antibody structure generation

Where we did not use experimentally determined antibody structures, structures were generated using the following tools: IgFold [10], ABodyBuilder2 [5], AlphaFold2-Multimer v2.2 [12], and ABlooper [20], all using default settings. For antibodies modelled using ABodyBuilder2, we ran refinement using openMM 8.0 [23] on all four generated models (as opposed to just the top1 model) in order to generate an ensemble of antibody models as described below.

### C. AlphaFold2 antigen modelling

AlphaFold2 v.2.2 was used to build antigen models for docking. Multiple sequence alignments (MSAs) for each of the antigen targets were built using the complete genomic databases used in the standard AlphaFold2 pipeline [8]. The ‘monomer ptm’ pipeline was used to predict antigen structures for the 69 monomer antigens, while the ‘multimer’ pipeline was used to predict the structures for the 2 multi-chain antigens (7r89 and 7k7h). Note that the bound antigen in 7k7h [24] contains a peptide, which was not included in the AlphaFold2 predicted structure. Both pipelines were run with Amber relaxation. The maximum template date for the modeling step was set to the same cut-off that was used for generating the benchmark set (July 2021) and a maximum of 20 template hits were used for the MSA generation step.

### D. AlphaFold2 antigen-antibody complex modelling

Multimer models for the antibody-antigen complexes were generated using AlphaFold2-Multimer v.2.2. The ‘multimer’ pipeline was used to predict structures of the complete antigen-antibody complex by including the sequences of both antibody chains and the antigen chains in the same .fasta file. As above, the AlphaFold2 generated structures were relaxed with Amber and a maximum of 20 template hits, the same template date cut-off (July 2021) was used as for antigen modelling.

### E. Information-driven HADDOCK3 protocols

In this work we exploit the high degree of modularity provided by the new version of HADDOCK, HADDOCK3 (github.com/haddocking/haddock3, unpublished), to run several information-driven docking workflows on our data set of antibody-antigen complexes. These differ in the interface information provided to define the ambiguous interaction restraints (AIRs) to drive the docking (see Supplementary Fig. 1 for an example representation of the three scenarios):

#### Para-Epi

The AIRs are defined based on the experimentally determined interface residues on both the antibody and the antigen. Residues are considered to be part of the interface if they are within 4.5 Å heavy-atom distance of the interaction partner in the reference structure. This represents a highly accurate definition of the paratope and epitope, reflecting the amount of knowledge on the antibody-antigen interaction likely present during the later stages of a drug discovery project. It is important to remark that this type of restraints does not introduce any pairing information (unlikely to be present in a real-case scenario), but rather couples each interface residue in the antibody (resp. antigen) to all interface residues in the antigen (resp. antibody).

#### CDR-VagueEpi

In this scenario, the AIRs are defined based on the residues in the CDR loops according to the IMGT definition to the true interface residues on the antigen (those within 4.5 Å from the antibody) and in addition all surface neighbors of those within a 4 Å cutoff. This effectively generates a larger interface on the antibody in order to allow for a realistic level of contacts surrounding the epitope. In this scenario, AIRs are only defined from the antibody CDR residues (defined as active in HADDOCK terms) to the antigen residues (defined as passive in HADDOCK terms), meaning that any epitope residues not contacting the antibody would not be penalized [21]. This represents a lower information scenario more likely to occur during the early stages of a project or during a virtual screening campaign.

#### CDR-VagueEpi-AA

The interface residues are defined as in CDR-VagueEpi, but both sets of residues are set as active during the docking process (i.e. an antigen residue not making contacts with the antibody will be penalized).

For each of these three scenarios we run nine docking protocols, shown in Table I, using both the bound structure of the antigen taken from the crystal structure of the complex and the AlphaFold2-predicted antigen structure. Our default docking protocol (DDP) is composed of the following steps:

1. **Rigid body docking phase**, in which 48 models are generated. This is a very low number compared to the number of models generated in standard HADDOCK sampling (typically 1000 or 10000 models [4, 21, 25]).
2. **Flexible refinement step**, the most CPU intensive step of the workflow, in which interface residues are treated as semi-flexible (first side-chains, then both sidechains and backbone) in order to allow for some binding-induced conformational changes following a simulated annealing protocol in torsion angle space [4].
3. **Final short energy minimisation**
4. **Fraction of Common Contacts clustering** [26], which clusters models sharing at least 60% of their interface contacts. A minimum of 3 models are required to define a cluster.

**TABLE 1.** Docking protocols. The protocols denoted ABBE or ENS use an ensemble of antibody models as input, formed either by four ABodyBuilder2 generated structures (ABBE) or by one structure generated by each ABodyBuilder2, AlphaFold2, IgFold and ABlooper (ENS).

| Protocol name | Antibody origin | Protocol |
| --- | --- | --- |
| ABB | ABodyBuilder2 [5] | DDP |
| ABL | ABlooper [20] | DDP |
| IG | IgFold [10] | DDP |
| AF2 | AlphaFold2-Multimer [12] | DDP |
| ABBE | ABodyBuilder2 ensemble | DDP |
| ENS | ABB-ABL-IG-AF2 ensemble | DDP |
| ENSNOAF2 | ABB-ABL-IG ensemble | DDP |
| ENS-196-48 | ABB-ABL-IG-AF2 ensemble | DDP but sampling 196 models during rigid-body |
| ENS-196-CLT | ABB-ABL-IG-AF2 ensemble | DDP but sampling 196 models during rigid-body with FCC clustering after rigid-body |

Both single models and clusters are scored using the default HADDOCK scoring function which consists of a linear combination of intermolecular van der Waals and electrostatic energies using the OPLS [27] force field, an empirical desolvation energy term [28] and the restraint energy. As an additional comparison, we also perform bound docking (using the bound structure of both antibody and antigen as input) for each of the three scenarios.

### F. ZDOCK protocol

We run ZDOCK version 3.0.2 [29] on our data set in order to compare the results with those obtained with HADDOCK3. ZDOCK is a rigid-body docking software using a FFT-based algorithm to evenly sample the roto-translational space of the protein-protein interaction. Existing information on the interface residues can be used a posteriori to filter the obtained complexes [29]. The residues used for filtering were either the true interface residues as defined above (Para-Epi) or the looser definition (CDR-VagueEpi)

We perform 16 ZDOCK runs for each complex in our data set, varying the input antibody structure (ABodyBuilder2, ABlooper, AlphaFold2 and IgFold), input antigen (bound and AlphaFold2 predicted), and information scenario (Para-Epi and CDR-VagueEpi, note that ZDOCK does not distinguish between active and passive residues). For each docking protocol, we run ZDOCK with default settings (15 degree sampling). Of the 2000 obtained solutions, we analysed the top 48 for consistency with the HADDOCK3 protocols.

## III. RESULTS

In the following, we first discuss the accuracy of the predicted antibodies and antigens. Secondly, we show that AlphaFold2-Multimer cannot accurately predict the structure of most antibody-antigen complexes. Then, we focus on the docking performance of HADDOCK3, comparing it to ZDOCK and highlighting that the usage of different antigen structures and docking protocols is critical for the accuracy of the resulting models. Finally, we investigate how the measured and predicted accuracies of epitope and paratope regions are correlated to the docking success rate.

### A. Antibody structure accuracy

We evaluated the modeling performance by calculating the backbone RMSD of the heavy chain CDR 3 (CDR H3) after aligning the modelled antibody to the experimentally determined reference antibody using the framework backbone coordinates [5] using IMGT numbering [22]. All RMSD calculations in this study were performed on backbone atoms. ABodyBuilder2 and AlphaFold2-Multimer (AF2) performed the best out of the four tested tools, with median CDR H3 RMSD of 2.34 Å and 2.15 Å and mean RMSD of 2.98 *±* 2.30 Å and 2.66 *±* 1.96 Å respectively, while ABlooper and IgFold achieved median RMSD of 2.68 Å and 2.63 ^Å^ and mean RMSD of 3.26 *±* 1.87 Å and 3.08 *±* 1.75 Å respectively. This holds when considering backbone RMSD across the paratopes, with ABodyBuilder2 and AlphaFold2 achieving similar median (1.72 Å and 1.74 Å) and mean (2.03 Å and 1.87 Å) paratope RMSD.

Fig. 1 reports the distributions for CDR H3 and paratope RMSD for the four antibody modelling methods. Additionally, we report the values of the lowest RMSD model in the ensemble generated by the four ABodyBuilder2 models and the ensemble consisting of one model generated by each of IgFold, ABodyBuilder2, AlphaFold2-Multimer, and ABlooper models. Model ensembles often increase the likelihood of an accurate antibody model being passed to the docking algorithm.

**FIG. 1.**
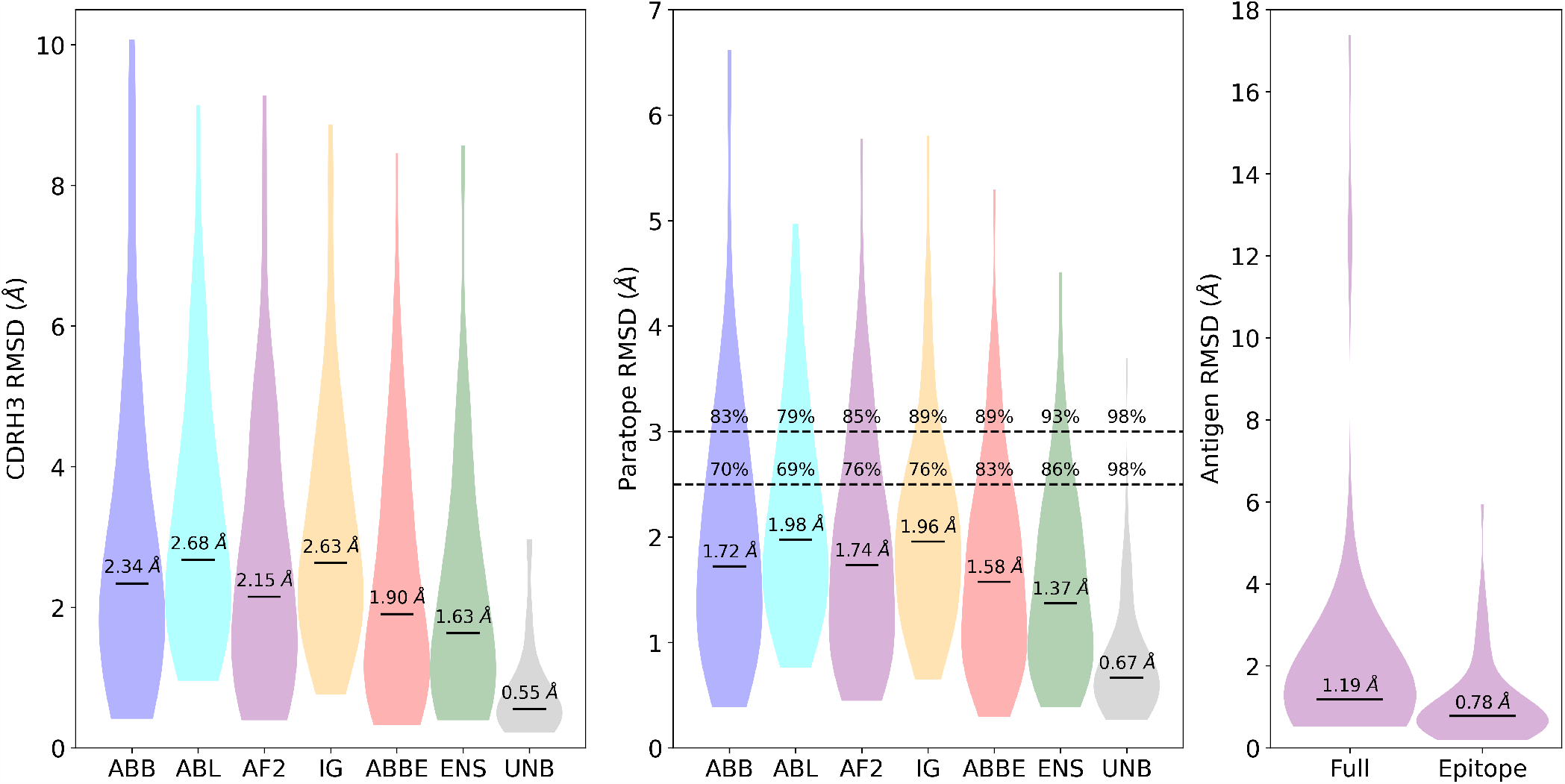
*Left* : violin plot of the distribution of backbone CDR H3 RMSD values for each of the four modeling methods (ABB, ABL, AF2, IG, see Tab. I), together with the value of the best model of the ABodyBuilder2 ensemble (ABBE) and of the ensemble of ABB, ABL, AF2, IG models (ENS). Values calculated over a set of experimentally determined unbound antibodies [30] (UNB) are reported for comparison. Highlighted values report the median CDR H3 RMSD for each considered method. *Center* : same but considering the backbone atoms RMSD of the amino acids that are part of the antibody paratope region. Besides the median values, this plot also contains the percentage of models that have paratope RMSD lower than 2.5 and 3.0 Å. *Right* : violin plot of the antigen accuracy expressed in terms of full structure and epitope RMSD with respect to the experimentally determined molecule.

### B. Antigen structure accuracy

We assessed the quality of the AlphaFold2 antigens by their backbone RMSD to the experimentally determined bound structure, both on the full backbone and epitope backbone only (see Fig. 1). The majority of antigen structures are modelled with high accuracy, with a median (resp. mean) overall RMSD of 1.19 Å (resp. 2.09 *±* 2.74 Å) and a median (resp. mean) epitope RMSD of 0.78 Å (resp. 1.10 *±* 1.00 Å). Of the 71 antigen structures, 50 are modelled with overall RMSD *<* 2.0 Å, 61 are modelled with epitope RMSD *<* 2.0 Å and only one antigen (7LFB) shows an epitope RMSD *>* 5.0 Å.

### C. Definition of docking success

To assess the performance of the various docking protocols used in this study, we quote the topN success rate (SR), that is, the fraction of structures in the data set that have at least one acceptable/medium/high quality complex in the top N ranked models according to CAPRI criteria [31, 32] (see Supplementary Material section 3). In the following we refer to the Top1/10 SR for acceptable/medium/high quality as T1/10-acc/med/high.

### D. AlphaFold2-Multimer performance

We assessed the performance of AlphaFold2-Multimer v2.2 on the data set as a baseline of a state-of-the-art machine-learning driven, unguided complex prediction algorithm for antibody-antigen modelling. AlphaFold2-Multimer achieved T1-acc of 0.17, T10-acc of 0.21, T1-med of 0.10, T10-med of 0.11, T1-high of 0.06 and T10-high of 0.07.

These results, comparable to those reported by Yin *et al*. [13] for antibody-antigen complexes, show that even though AlphaFold2-Multimer reaches notable results in predicting protein-protein complexes [12], the accurate modelling of antibody-antigen complexes is still challenging and could benefit from classical physics-based methods, as shown below, especially when some information is available to drive the modelling.

### E. Rigid-body docking modelled antibodies against bound antigens achieves high success rates

We next analysed the performance of the data-driven docking scenarios listed in Table I using only the comparatively computationally inexpensive rigid-body docking stage of HADDOCK and experimentally determined bound antigen structures. We compared the influence of both different levels of information on the paratope and epitope location (see Methods) as well as the choice of the antibody ML modelling tool. We further established an upper bound of performance by docking the experimentally determined antibody structure against the experimentally determined antigen structure (bound-bound docking).

As has been previously shown by Ambrosetti *et al*. [4], precise information on the paratope and epitope (the Para-Epi scenario) results in accurately generated binding poses, with T1-acc of 1.0 for the bound-bound baseline. The docking performance using computationally modelled antibodies is dependent on the modelling tool used. The two non-ensemble protocols with the lowest median CDR H3 RMSD on the data set (ABB and AF2) performed best after the rigid-body docking stage, with a T1-acc of 0.56 and 0.56 and a T10-acc of 0.90 and 0.93 respectively (Fig. 2). Using a level of information on the paratope and epitope residues as could be expected in a virtual screening scenario [33] to drive docking (the CDR-VagueEpi scenario, see Methods) reduces the performance for all methods (see Fig. 2). This was expected given the lower quality of the available information and has been demonstrated previously [4] on experimentally determined antibody structures. We found that the CDR-VagueEpi-AA protocol outperforms the CDR-VagueEpi protocol in most tested scenarios, with the best non-ensemble input protocol by T1-acc (ABB) achieving T1-acc of 0.37 and T10-acc of 0.58 for the CDR-VagueEpi-AA protocol, compared to 0.25 and 0.51 for the CDR-VagueEpi protocol. This trend holds for T1/10-med and T1/10-high. We therefore excluded the CDR-VagueEpi protocol from further analysis.

**FIG. 2.**
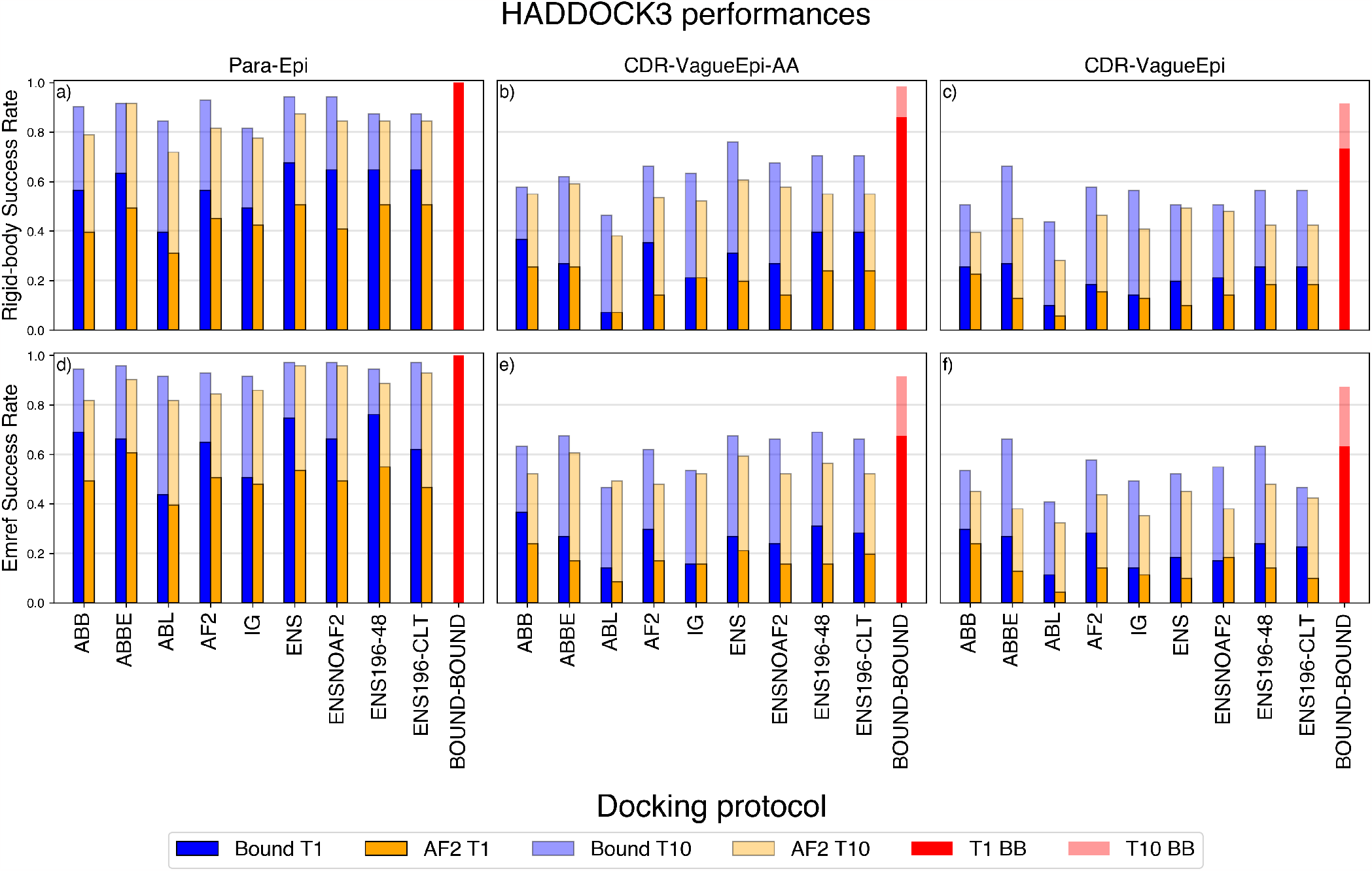
Success rate in the top 1 and top 10 poses using CAPRI criteria for defining an acceptable model. Dark blue bars (BOUND T1) refer to the SR of the top one structure for the bound antigen, while dark green bars (AF2 T1) concern the AF2-modelled antigen. Light blue (resp. light green) refer to the success rate (SR) over the top 10 poses for the bound antigen (resp. AF2-modelled antigen) scenario. *Top*: success rates after the rigid-body docking phase. *Bottom*: success rates after flexible refinement and energy minimisation. Protocols are briefly described in Tab. I.

### F. Using ensembles of models improves the success rate

We further investigated the effect of introducing structural diversity into the docking process by using ensembles of modelled antibody structures as input. Considering multiple computationally generated antibody structures in immunoinformatics pipelines has been previously identified as an important factor in antibody property prediction [34, 35]. We used two different approaches to the generation of model ensembles: distinct models were either generated using different modelling tools (ENS protocols) or using several structures generated by the same tool (ABBE protocol), see Table I. We further varied the number of distinct rigid-body poses generated in the first step of the HADDOCK protocol to either be the same as in the non-ensemble protocols (keeping the computational cost of that step equivalent to the non-ensemble protocols, but sampling each input antibody structure less) or generating as many rigid-body poses for each input model as were generated in the non-ensemble protocols, increasing the total number of rigid-body poses generated proportionally to the number of poses in the ensemble (see Table I).

We found that ensemble approaches result in improved success rates in the rigid-body step for the Para-Epi knowledge scenario (Fig. 2(a-c)), with the best ensemble approaches on Para-Epi by T1-acc (ENS, ENSNOAF2) achieving T1-acc of 0.68 (0.65 for ENSNOAF2) and T10acc of 0.94, compared to 0.56 and 0.93/0.90 respectively for the best single-input approaches (AF2, ABB). On the CDR-VagueEpi-AA scenario, the best single-input method by T1-acc (ABB) performed similarly to the best ensemble methods (ENS196-48/ENS196-CLT) on T1-acc, but the ensemble methods performed better on T10-acc and at T1/10-med/high, with a T1-med and T10-med of 0.18 and 0.34 for ABB and T1-med and T10med of 0.23 and 0.37 for ENS-196-48 (see Supplementary Fig.3-4).

The computationally least expensive ensemble approach (ABBE, using the output of only one structural modelling tool and sampling the same amount of rigidbody poses as the single-input approach) performs comparable to computationally more expensive approaches using output from several modelling tools and/or sampling more rigid-body poses.

### G. Using AlphaFold2 modelled antigens reduces the success rates

In the previous sections we reported the results using experimentally determined bound antigen structures. However, as modern computational pipelines often rely on computational modelling of antigens as well as antibodies, we analysed the impact of using AlphaFold2 antigen models on the docking performance. As expected, the performance drops when compared to the experimentally determined antigen structures across all knowledge scenarios (see Fig 2(a-c)). After the rigid-body stage, the best T1-acc using the Para-Epi scenario dropped from 0.68 for the ensemble models (ENS) using the experimentally determined antigen structure to 0.51 using the modelled antigen structure and the best T10-acc correspondingly dropped from 0.94 to 0.87. These trends hold for the less accurate knowledge scenarios (see Fig.2(b-c)) and the generation of med/high quality poses (see Supplementary material section 2), with ABBE being the most reliable workflow using modelled antigens.

### H. Performance of HADDOCK refinement protocol is scenario dependant

We further examined the impact of the HADDOCK refinement step on docking pose quality. In the Para-Epi knowledge scenario, where accurate information on the paratope and epitope residues is provided, introducing flexibility in the models (Fig. 2(d)) is beneficial to the docking success rate with an average improvement of T1-acc of 0.05 for protocols using the bound antigen and of 0.06 when considering the AlphaFold2-modelled antigen. The largest improvement can be observed in T1/10-med (as opposed to the T1/10-acc) (see Supplementary Fig. 3). Interestingly, for the CDR-VagueEpi-AA scenario, refining the rigid body models (Fig. 2(e-f)) has less effect on the success rate for both experimentally determined (average T10-acc change -0.02) and AlphaFold2-modelled antigens (average T10-acc change 0.0), in some cases even reducing it. These results indicate that while the refinement step consistently improves the success rate for the Para-Epi scenario, it is much less reliable in the CDR-VagueEpi-AA scenario. In such a setting, the computationally much less expensive rigidbody docking step can be sufficient for downstream use of the docking poses.

**FIG. 3.**
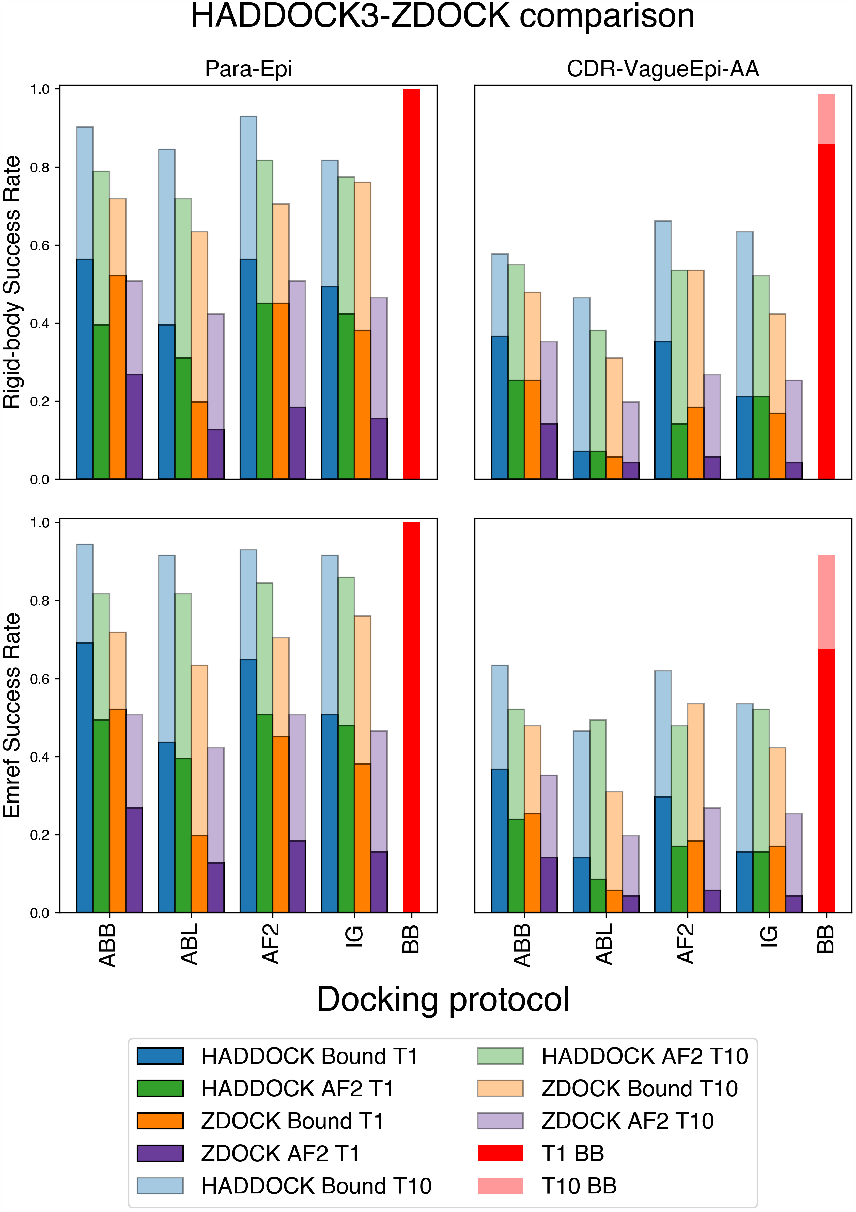
Comparison between HADDOCK and ZDOCK success rates in the top 1 and top 10 poses. Dark blue and dark green bars (HADDOCK BOUND T1, HADDOCK AF2 T1) refer to the SR of the top one structure for the bound and AF2-modelled antigen respectively. Light blue and light green bars (HADDOCK BOUND T10, HADDOCK AF2 T10) refer to the SR over the top 10 poses for these two scenarios. Orange (ZDOCK BOUND T1, ZDOCK BOUND T10) bars indicate the accuracy of ZDOCK runs using the bound antigen, while purple bars (ZDOCK AF2 T1, ZDOCK AF2 T10) correspond to ZDOCK runs targeting the AF2-modelled antigen. Finally red bars report the success rates of HADDOCK3 bound-bound docking as a baseline. *Top*: success rates after the rigidbody docking phase. *Bottom*: success rates after flexible refinement and energy minimisation. Protocols are briefly described in Tab. I.

We also show that applying a short energy minimisation to the models produced at the rigid-body docking stage slightly increases the T1-acc success rate, with an average improvement of 0.043 (resp.0.024) in the Para-Epi (resp. CDR-VagueEpi-AA) scenario (see Supplementary Material section 6).

A comparison between CPU usage required by the different HADDOCK protocols is available in the Supplementary Material (section 7).

### I. The HADDOCK protocol outperforms the ZDOCK rigid-body baseline

We compared the performance of HADDOCK3 and ZDOCK (see Methods). Fig. 3 shows the success rates over the two different scenarios (Para-Epi and CDRVagueEpi-AA), comparing ZDOCK to the HADDOCK3 models obtained both after rigid body (Fig. 3 (a-b)) and after refinement and energy minimisation (Fig. 3 (c-d)). HADDOCK3 outperforms ZDOCK on all scenarios and docking protocols. When considering docking runs with an AlphaFold2-modelled antigen, the chances of producing an acceptable pose with HADDOCK3 are on average two times higher than with ZDOCK.

### J. Paratope and epitope model accuracy are strong predictors of success rate

The success of protein-protein docking experiments strongly depends on how close the input structures are to their bound analogue [36, 37], which is typically unknown in a realistic scenario. It is quite intuitive to imagine that a docking protocol in which a badly modelled antibody is docked against a badly modelled antigen will not produce any acceptable model, irrespective of the quality of the information (restraints, see Sec. II E) used to drive the docking. Here we aim to inspect the extent to which this applies to the docking of ML-derived antibodies and antigens. To this end, we calculated the RMSD between the residues in the modelled epitope and paratope and the true bound structures. We do this by aligning the modelled and true paratope and epitope independently onto the reference complex. The resulting RMSD corresponds to the minimum interface-RMSD (min i-RMSD) achievable using the modelled structures with perfect rigidbody docking. Fig. 4 relates this quantity to the maximum DockQ [38] score obtained in the top 10 docking poses (DockQ-T10) for both the Para-Epi and the CDRVagueEpi-AA scenario. We observe a moderate negative correlation in the CDR-VagueEpi-AA scenario for all models between i-RMSD and DockQ-T10, with the Pearson correlation coefficients ranging between *ρ* = *−*0.29 for ABlooper to *ρ* = *−*0.51 for ABodyBuilder2. For all models, this correlation is stronger in the Para-Epi scenario, ranging from *ρ* = *−*0.58 for ABlooper to *ρ* = *−*0.64 for IgFold.

**FIG. 4.**
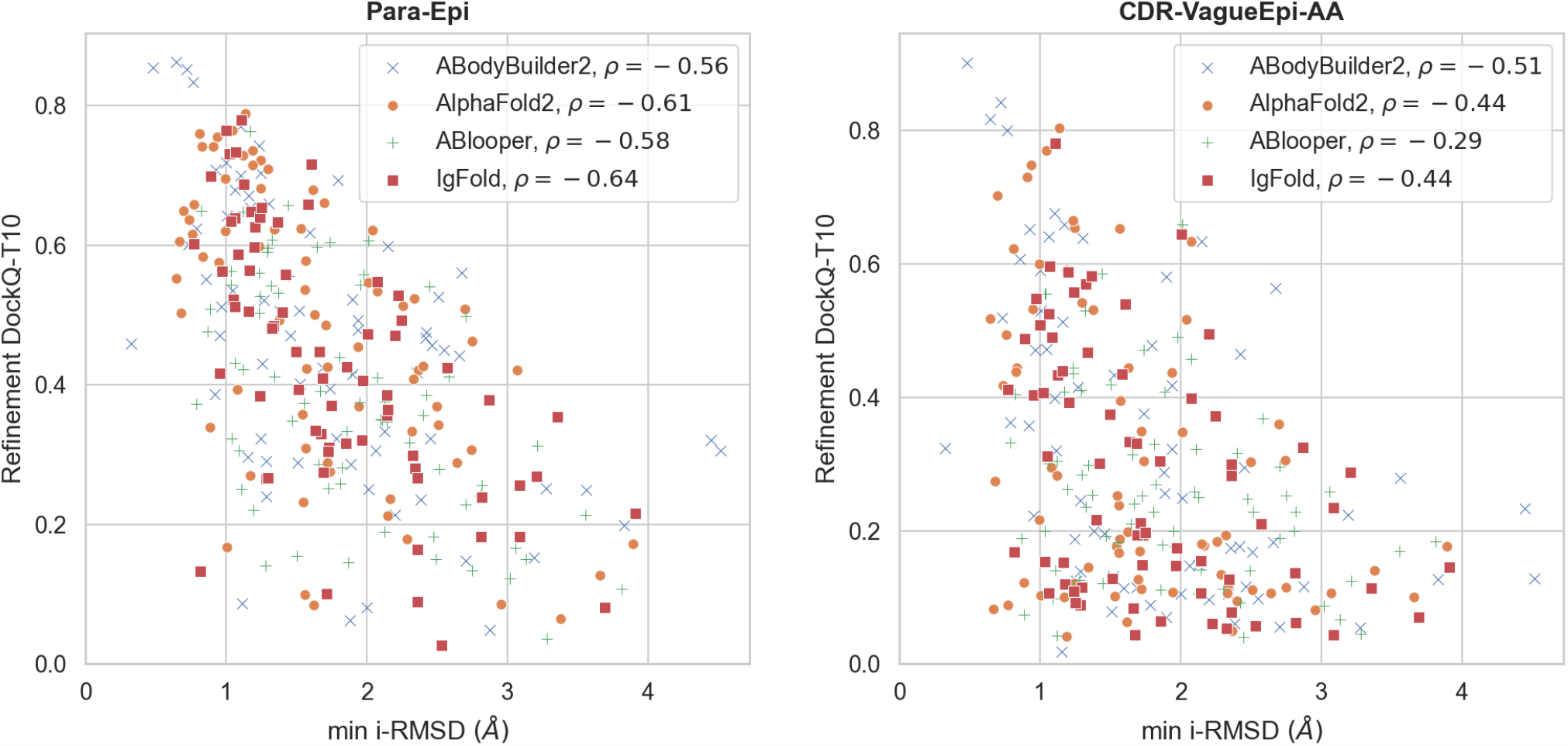
Maximum DockQ score in the top 10 ranked poses (DockQ-T10) against min i-RMSD after refinement. The left panel shows the Para-Epi scenario and the right panel shows CDR-VagueEpi-AA. Each point represents a different complex, with different colors/shapes indicating different antibody prediction models as described in the legend. In the legend we also list the Pearson correlation coefficient *ρ* for each data set.

We also observe that the accuracy of paratope residue modelling is more strongly correlated with docking success than the epitope modelling accuracy (see Supplementary Material section 4).

### K. Relationship between AlphaFold2 plDDT, ABodyBuilder2 confidence score, and docking success rate

The results shown above suggest that combining HADDOCK3 with ML-based structure prediction methods can yield reasonably accurate complex structure predictions, especially when the input structures are modelled with high accuracy. However, predicted structures are typically only used when there is no experimental structure available, and as such the model accuracy is not accessible. Many state-of-the-art protein structure prediction methods provide a per-residue confidence metric [5, 8]. Here we investigate the extent to which the average confidence scores of epitope and paratope regions of the predicted antibodies and antigens correlate with the accuracy of the models and the docking success rates. We report results for three confidence metrics: the AlphaFold2 average predicted local distance difference test ((pLDDT [8, 39]), averaged separately over paratope and epitope residues, and the ABodyBuilder2 root mean squared predicted error (RMSPE, [5]), averaged over the paratope residues.

The average epitope pLDDT is strongly inversely correlated with epitope RMSD (*ρ* = *−*0.65, see Supplementary Fig. 8), indicating it is giving a good measure of the epitope prediction quality. In terms of docking performance, the correlations with DockQ-Top10 differ by scenario, with correlation coefficients ranging between *ρ* = 0.10 (ABL) and *ρ* = 0.15 (ABB) for the CDR-VagueEpi-AA scenario and *ρ* = 0.17 (ABL) and *ρ* = 0.40 (ABB) for the Para-Epi scenario (see Supplementary Fig. 12). Similarly, the average paratope pLDDT shows strong anticorrelation with the paratope RMSD (*ρ* = *−*0.68, see Supplementary Fig. 8). Average paratope pLDDT correlates to DockQ-T10 for AlphaFold2 modelled antibodies, with *ρ* = 0.15 and *ρ* = 0.42 in the CDR-VagueEpi-AA and Para-Epi scenarios (see Supplementary Fig. 14).

ABodyBuilder2 paratope RMSPE is strongly correlated with paratope RMSD (*ρ* = 0.72, see Supplementary Fig. 10) and moderately correlated with DockQT10 in the CDR-VagueEpi-AA (*ρ* = *−*0.35) and Para-Epi (*ρ* = *−*0.45) scenarios (see Supplementary Fig. 16).

In summary, both plDDT and RMSPE show correlation with docking success and enable filtering of badly docked complexes in the Para-Epi scenario. For a more detailed breakdown of these results, see Supplementary Material section 4.

## IV. DISCUSSION

Predicting the complex structures of antibody-antigen complexes is one of the fundamental tasks within computational antibody discovery. In this study, we have established a benchmark data set for the evaluation of ML-driven modelling of this class of complexes. By ensuring that the benchmark data set has no overlap with training sets for current ML-driven modelling tools, we have generated a large and diverse data set that can be used to objectively assess current state-of-the-art approaches for antibody-antigen modelling.

Modern antibody-antigen modelling workflows rely increasingly on ML-generated input structures of both the antibody and the antigen. In this study, we have assessed the impact of using ML-generated antibody and antigen structures on the performance of different information-driven protocols using HADDOCK. While there is clearly a reduction in performance from using ML-generated input structures compared to bound experimentally determined structures, useful interface models can be generated for most cases. We explored a low information scenario where the paratope and epitope residues were specified coarsely, together with ML-generated antibody and antigen structures. This corresponds to the typical data available during the early stages of a computational antibody design campaign.

In doing so, we also demonstrated that flexible refinement of poses derived from ML-modelled antibodies and antigens can improve the docking performance in some scenarios, but that computationally fast rigidbody docking alone already achieves good results. We further showed that utilising the ability of ML-driven antibody modelling tools to generate ensembles of input structures markedly improves the docking performance and thereby highlighted a computationally efficient way to boost the performance of antibody-antigen docking protocols in general. The protocol using an ensemble of models from ABodyBuilder2 (ABBE) and AlphaFold2 generated antigen structures achieves a high success rate in predicting acceptable complexes at rather low computational costs, outperformed our baseline (the ZDOCK rigid-body docking program) and other single-tool protocols tested, achieving performances comparable to the ensemble protocols.

Unsurprisingly, the docking success is highly dependent on the quality of the modelled structures, as expressed in terms of interface, paratope, and epitope RMSD, calculated with respect to the experimental bound structures. The lower these values, the easier it is for the HADDOCK protocol to sample and to properly rank accurate models of antibody-antigen complexes. In a real scenario, the quality of the generated models will be unknown. Our analysis revealed that the confidence scores for antigens and antibodies, produced by AlphaFold2 and ABodyBuilder2 are predictive of the quality of the individual antibody and antigen models and are moderately correlated with docking success. This is particularly true when accurate information on the paratope and epitope is available (the Para-Epi scenario). Further work is warranted to improve the ability of these measures in predicting docking success, ideally including also the quality of the provided interface information for a better prediction of the reliability of the docked models.

This work, which has established a benchmarking data set and proposed a ML-driven, ensemble docking approach clearly highlights the potential of ML-driven docking approaches for the high-throughput analysis of antibody-antigen interactions.

## DATA AVAILABILITY STATEMENT

The code to generate and analyse the data is freely available at github.com/haddocking/ai-antibodies. The docking models generated in this study will be deposited at data.sbgrid.org/labs/32/ upon acceptance.

## VI. AUTHOR CONTRIBUTIONS STATEMENT

A.M.J.J.B and C.D. supervised the project. C.S. and N.D. prepared the benchmarking data set. M.G. developed the docking protocols and performed the calculations. M.G., C.S., and D.C analysed the data. M.G., C.S., D.C, and N.D. wrote the manuscript. All authors reviewed the manuscript.

## VII. ACKNOWLEDGMENTS

This project has received funding from the European Union Horizon 2020, projects BioExcel (823830 and 101093290) and EGI-ACE (101017567), and from the Netherlands e-Science Center (027.020.G13).

