## Supplementary Material for "Towards the accurate modelling of antibody-antigen complexes from sequence using machine learning and information-driven docking"

### Supplementary Material for
### "Towards the accurate modeling of antibody-antigen complexes from sequence using machine learning and data-driven docking"

Marco Giulini[1], Constantin Schneider[2], Daniel Cutting[2], Nikita Desai[2],
Charlotte M. Deane[2], Alexandre M.J.J. Bonvin[1]

[1]Bijvoet Centre for Biomolecular Research, Faculty of Science - Chemistry, Utrecht University, Padualaan 8, 3584 Utrecht, CH, The Netherlands
[2]Exscientia plc, The Schroedinger Building, OX4 4GE, Oxford, UK

November 17, 2023

#### Contents

**1 Data set** **3**

**2 On the effect of clustering on docking success rate** **5**

**3 An analysis of medium and high-quality docking poses** **6**

**4 Influence of epitope and paratope RMSD on docking quality** **9**

**5 Epitope and paratope confidence scores as predictors of docking quality** **14**

**6 Energy minimisation of rigid-body solutions increases the top 1 success rate** **21**

**7 Protocol timings** **22**

**References** **23**

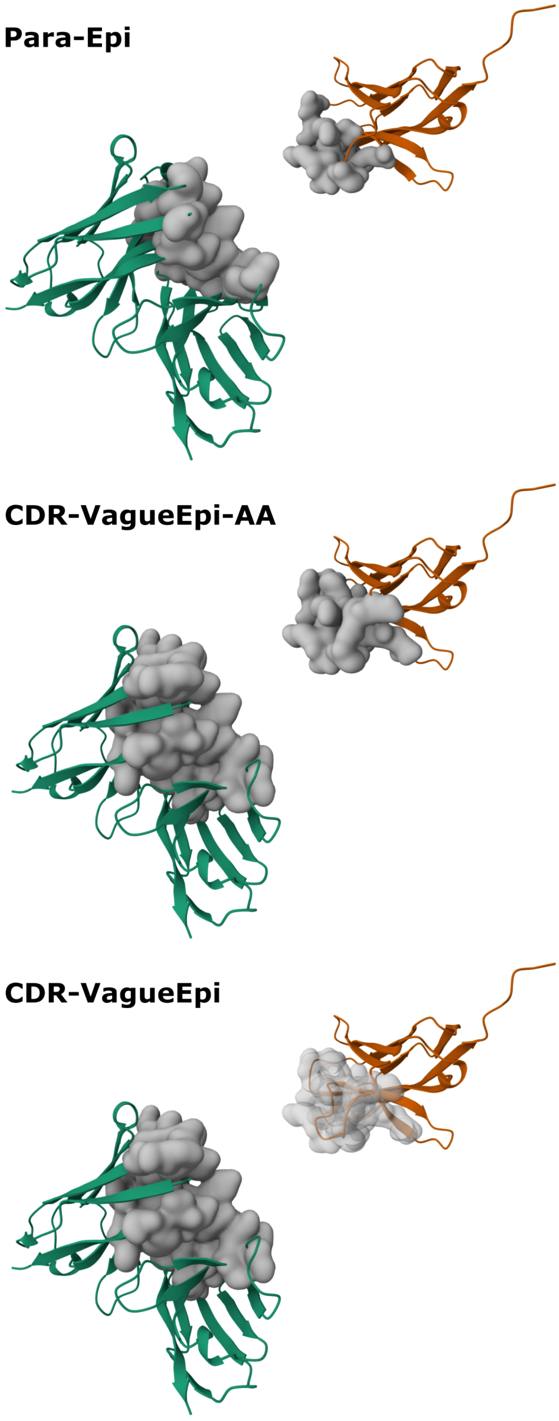


**Supplementary Figure 1:** Representation of the regions defined as important for the interaction (i.e., converted into ambiguous interaction restraints) in the three scenarios for pdb 7VUX. The (bound) antibody structure is represented in green, while the antigen is coloured in orange. The **Para-Epi** scenario contains information about the true interface, while in the two **CDR-VagueEpi** scenarios all the CDR loops are labeled as potentially active. The only difference between these two scenarios lies in how we treat the vague definition of the epitope for the antigen (labelled as active in **CDR-VagueEpi-AA** and as passive in **CDR-VagueEpi**).

### 1 Data set

Supplementary Table 1: CDR loop sequences of the 71 antibodies present in the data set

| PDB | H1 loop | H2 loop | H3 loop | L1 loop | L2 loop | L3 loop |
|---|---|---|---|---|---|---|
| 7bbj | ASGYTFTTYWMH | GAIYPGLSDTTY | YYCARLLDYAMDYW | CRASQDIRSY | LLIWYTS | ATYFCQQGETLPW |
| 7bnv | GSGYSFTSYWIG | GIIYPGDSDTRY | YYCARHPSPYYYGSGSYSGGFDYW | CRASQSVSSSY | LLIYGAS | AVYYCQQYDNWPLMH |
| 7daa | VSGFSLSDYAMS | GIIYASGSTYY | YFCARYYAGSDIW | CQASQSISAY | LLIYDAS | ATYYCQTYYAIITYGA |
| 7dk2 | ASGFTFSSYWMS | ANIKQDASEKYY | YYCARDLGILWFGDYPW | CRASQGISNS | LLLYAAS | ATYYCQQFYSTPR |
| 7e3o | ASGFTFSSYAMH | ALISYDGSNKY | YYCARGLGLRFLEWPISSYW | CSGSSSNIGSNT | LLIFSNN | ADYYCAAWDDSLKG |
| 7e5o | VYGGSFSGYYWS | GEINHSGSTNY | YYCARGWDLDYYYGMDVW | CRASQGIGNY | RLIYAAS | ATYYCLQHGFLPW |
| 7e72 | ASGYSFTSYWMN | GMIHPSDSETRL | YYCARGLYGNSW | CRASQDIGIS | RLIYATS | VDYYCLQYASSPY |
| 7f7e | ASGFTFSSYAMS | SAIVGSGGSTYY | YYCAKSLIYGHYDILTGAYYFDYW | CRASQGIGNW | LLIYAAS | ATYYCQQANSFP |
| 7k7h | ASGYAFTNYKAL | GYIDPYNSDSSY | YYCAGLELTGTLPYW | CKSSQSLFNSRTRKNH | LMIYWAS | AVYLCKQSHNRAL |
| 7k9j | ASGYTFTSYWIT | GDIYPGSGSTKY | YYCARWDFYGSRTFDYW | CRASQNIGTI | LLIKYAS | ADYYCQQSSSWPL |
| 7kez | ASGFNIKDTYIH | ARIYPTNGYTRY | YYCARGGAVAGTGVYYFDYW | CRASQDIPRSISGY | LLIYWGS | ATYYCQQHYTTPP |
| 7kf0 | ASGFNIKDTYIH | ARIYPTNGYTRY | YYCARGGSFYYYYMDVW | CRASQDIPRSISGY | LLIYWGS | ATYYCQQHYTTPP |
| 7kf1 | ASGFNIKDTYIH | ARIYPTNGYTRY | YYCAKLGIGYYYYGMDVW | CRASQDIPRSISGY | LLIYWGS | ATYYCQQHYTTPP |
| 7kn3 | ASGFTVSLSHMN | SITYGDGNSDY | YYCAREYYYGMDVW | CKSSQSVLYSSNNKNY | LLIYWAS | AVYYCQQYYSLPL |
| 7kql | VSGGSISSRSYYWG | GSIYYSGFTY | YYCATGGPYGDYAHWFEPW | CRASQSVSSSY | LLIYGAS | AVYYCQQYGSSPI |
| 7kyo | ASGITFSSYAMS | ASISSGGSTYY | YYCARGPMALLYYRGFDYW | CKASQSVDYDGDSY | LLIYAAS | ATYYCQQSNEDPW |
| 7l7r | ASGFTFSSYVMS | SVIYRGGSTKY | YYCVKDPKAWLEPEWW | CRASQSISKY | LLIYAAS | ATYYCQQSYSNPR |
| 7lf7 | ASGFTFSRCAMS | SAISRDSTYTYY | YYCARQIDDYYVDALDYW | CRASKIISKY | LLIYSGF | AMYYCQQHNEYPL |
| 7lfa | ASGYTFTSHWMQ | GAIYPGDGDTKF | YYCARENLYGYYFDYW | CRSSTGAVTSGNF | GLIGGAD | AIYFCALWYSDHW |
| 7lfb | TSGYAFTNYGVN | GWINTNTGQTTY | YFCARLIYDGDYISSDFW | CGASENIYGA | LLIYGAT | ATYYCQNALSMPY |
| 7lr3 | ASEFTFSDYGMH | ASISSGSNSIYY | YYCSREAYFAMDYW | CKASQDIHKY | LLIYYTS | ATYFCLQYDNLY |
| 7lr4 | ASDYSLSDYNMN | GVINPNHGTTHY | YYCASPIHYGNHVPFDYW | CRTGQDISNY | LLIYFTS | ATYFCQQGITLPW |
| 7mdj | ASGYTFTSDWIH | GEIIPSYGRANY | YYCARERGDGYFDYW | CRASQSIGTD | LLIKYAS | ATYYCQQSNRWPF |
| 7mrz | VSGGSISSSSYYWG | GSISYSGSTYY | YYCARDSLRYGMDVW | CKSSQSVLYSSNNKNY | LLIYWAS | AVYYCQQYALAPPR |
| 7msq | VSGGSISSYHWN | GYIYYSGNTNY | YYCVREMRRGYSGYDYWDLYAFDIW | CRASQGISSY | LLIYAAS | ATYYCQQLNSYPH |
| 7mzf | ASEFIVSRNYMS | SVIYSGGTTYY | YYCARDRGDYLFDYW | CRASQSISSW | LLIYKAS | ATYYCQQYNSYFP |
| 7mzg | ASGLTVSSNYMS | SVFYPGGSTYN | YYCARDAVYYGMDVW | CRASQSISSY | LLIYAAS | AIYYCQESYSTPGLF |
| 7mzh | ASGYTFTGYYMH | GRINPNSGGTNY | YYCARSYYDYW | CSGSSSNIGHNA | LLIYYDD | ADYYCAAWDDILNGP |
| 7mzi | ASGFTFSRFAMT | SAISGSGGSTYY | YYCAKVGWGAFDIW | CSGTYSNIGSNP | LLIYAND | ADYYCSTWDDSLPGP |
| 7mzj | ASGFTFSYAWMS | GRIKRKSDGGTTD | YYCTTDLCRSTSCEHDAFDIW | CRASQSIRSY | LLIYAAS | ATYYCQQSYTTPAI |
| 7mzk | ASGYTFTSYYMH | GIINPSGGGTSY | YYCAKDRVTIFWGNGMDVW | CKSSQSVLYSSNNKNY | LLIYWAS | AVYYCHQYSTTPL |
| 7n0u | VSGGSITNYFWT | GYIYYSGGTNY | YYCAGSYYYGVDVW | CRASQSIKSF | LLIYDAS | AVYFCQQRNNWPF |
| 7n3i | ASGFTVSSNYMS | SVIYSGGSTYY | YYCARDLYSSGGTDIW | CRASQSVSSSY | LLIYGAS | AVYYCQQYGSSPG |
| 7n4i | GSGYSFISYWIA | GIIYPGDSDTTY | YYCARLLYYSDSSPLDSW | CRASQSISTY | LLIYAAS | ATYYCQQSHSTPR |
| 7n4j | VSGDSISSSDYSWG | GTIYYIKNTY | YYCARERPPFDVVVVPAARPYNWFDP | CTGSSSNIGAGYD | LLIYGN | ADYYCQSYDSSLSGSK |
| 7np1 | ASGITVSSNYMS | SVIYSGGSTYY | YYCARGEGGSIVGVTSDYW | CRASQSISRY | LLIYAAS | ATYYCQQSYSTLPY |
| 7nx3 | ASGYAFSSYWVN | GQIYPGDGDTNY | YFCARSRGYFYGSTYDSW | CRASESVDNYGISF | LLIYAAS | AMYFCQQSKEVPW |
| 7phu | ASGYTFTDYYIH | GWINPNSGGTNY | YYCARDLWFGESPPYYGVDVW | CGGNNIGSYS | LVIYYDS | ADYFCQVWDTNTDHV |
| 7phw | ASGFDFSSYAMG | AGIRNDGSFTLY | YFCTKSADDGGHYSDFSGEIDAW | CSGSTYN | TVIYYND | AVYYCGNSDSRNV |
| 7pi7 | ASGFNIKDTYIH | GRIDPANGNTYS | YYCARDVLYFDVW | CRASESVDSYGNSF | LLISRAS | ATYYCQQSNEDR |
| 7pqy | ASGFTVSSNYMS | SVIYSGGSTYY | YYCARDHVRPGMNIW | CQASQDISNY | LLIYDAS | ATYYCQQYDNLPV |
| 7pr0 | ASGFTFSSYSMN | SYISSSSSTIYY | YYCASPGGITAAGTSVLFGYYGMDVW | CRSSQSLLHSNGYNY | LLIYLG | GVYYCMQALQTPITW |
| 7ps0 | VSDGSISSSDYYWS | GYIYYTGSTYY | YYCARLVVPSPKGSWFDPW | CTGTSIDVGNYNL | LIIYEGS | ADYYCCSYVGSSTY |
| 7ps1 | ASGLTVRSNYMN | SLIYSGGSTFY | YYCARDLVVYGMDVW | CRASQSVSSSS | LLIYGTS | AVYYCQQYGSSP |
| 7ps2 | ASGFTFSNYGMH | ALISYEESNRYY | YYCAKDQGPATVMVTAIRGAMDVW | CKSSQSVLYSSNNKNY | LLIYWA | AVYYCQQYFGSPSI |
| 7ps4 | GSGYSFTNYWIG | GIIYPGDSGTRY | YYCARSRVGATGGYYDYYMDVW | CSGSSSNLGGNT | LLIYSNN | ADYYCAAWDDSLNGP |
| 7ps6 | VFGGSITSSNHYWV | GSMYYSGSTA | YYCARQIGPKRPSQVADWFDPW | CRASQGISSY | LLIYAAS | ATYYCQQLNSYPL |
| 7q0g | ASGGTFSSSVIS | GGIIPLFGSANY | YYCAKVSQWALILFW | CRASQSVSSSY | LLIYGAS | AVYYCQQYGTSPSW |
| 7q0i | ASGFTFSSYGMH | AVIWYDGSNNFY | YYCARSYCSGGFCFGYYYGLDVW | CGGNNIGTKS | LVIYYNS | ADYYCQVWDSGSDHY |
| 7qnw | VSGDSISSSRYYWG | GTFYYSGITY | YYCARPRPPDYYDNSGALLFDIW | CRASQSISAW | LLIYKAS | ATYYCQQYISSSPW |

Continued on next page

Supplementary Table 1: CDR loop sequences of the 71 antibodies present in the data set (Continued)

| | | | | | | |
|---|---|---|---|---|---|---|
| 7qny | ASGFTFDDYAMH | SGVSWNSGTIGY | YYCAREVGGTFGVLISREGGLDYW | CGGNTIGSKS | LVVYDDS | ADYYCQVWDSSSDRV |
| 7qu1 | ASGYAFGSHWMN | GQIYPGDGDTNY | YFCARDDYGTRYYFDYW | CRASQDINNY | LLIHYTS | ATYFCQQGKTLPL |
| 7qu2 | TSGFTFSNYQMH | AVITVKSDNYGAN | YFCSRSGIYDGYYAYAMDYW | CKASQIVGTS | LLIYWAS | ADYFCQQYATYPL |
| 7r89 | TSGFTFSEFFME | AVSRNEANDYTTD | YYCARDAWMGFDYW | CRASQEISGY | RLIYAAF | AHYYCLQYASYPC |
| 7r8l | ASGITVSSNYMS | SVMYAGGSTFY | YYCARDLYSSGGTDIW | CRASQSIGSSY | LLIYGAS | AVYYCQQYGSSPG |
| 7rah | ASGFTFSSYGMS | ATISSGGTYTYY | YYCAREIMRGGGYYFDYW | CRASQDISNY | LLIYYTS | ATYFCQQGNTLPY |
| 7rco | VSGFSLSSYTVN | GYISYGGSAYY | YFCARHMQVGGAPTGSMAAFDPW | CQSSQSVYNNNY | LLIYGAS | ATYYCAGGYSGSSDKY |
| 7rfb | TSGGTVNTLH | GSIFPLLGVPTY | YYCAKDGVGWSGHGPSQWSGVDVW | CTSSQSLLHSTGYNY | LLIYLG | GIYYCMQALEIPRL |
| 7rks | ASGFTFSNYAMH | AVISYDGSNKYY | YYCASGYTGYDYFVRGDYYGLDVW | CTLSSGHSSYA | YLMKLNTDGS | ADYYCQTWGTGIL |
| 7s0b | ASGFTFSSYAMS | SLISGSGGSTYY | YYCARDLWGSGFFAFDVW | CQASQDISNY | LLIYDAS | ATYYCQQDAGTPL |
| 7s11 | VSGLSLTTNSVS | GVIWSNGGTDY | YFCARNFPYPGINFDW | CHSSTGAVTTSNY | AILGGTS | ATYFCSLWYSGHL |
| 7s13 | VSGLSLTNNIVS | GVIWSNGGTDY | YFCASRDYPGFAYW | CVGDELPKRY | RVIYEDS | ADYYCLSTYSDDKLP |
| 7s4s | ASGFIVSRNYMI | SVIYSGGSTFY | YYCARDLEVAGAIDYW | CRASQSISSSY | LLIYGAT | AVYYCQQYGSSPGY |
| 7seg | ASGYTFTSYYMH | GAIEPMYGSTSY | YYCARGSAYYYDFADYW | CGGHNIGSKN | LVIYQDN | ADYYCQVWDNYSV |
| 7sem | ASGFTFSSYSMN | SSISASSSYSDY | YFCARARATGYSSITPYFDIW | CTGSSSNIGAGYD | LLIYDNN | ADYYCQSYDRSLSG |
| 7shu | VSGYNITSGYSWN | ASVTYDGSTNY | YYCAKGNNYFGHWHFAVW | CRASKSVDSDGDSY | LLIYAAS | ATYYCQQSHEDPY |
| 7shz | VSGYSITSGYSWN | ASIKYSGETKY | YYCARGSHYFGHWHFAVW | CRASKPVDGEGDSY | LLIYAAS | ATYYCQQSHEDPY |
| 7si0 | VSGYSITSGYSWN | ASVTYDGSTNY | YYCARGSHYFGHWHFAVW | CRASQSVDSDGDSY | LLIYAAS | ATYYCQQSHEDPY |
| 7so9 | ASGFTFNSYGMH | AFIRYDGGNKYY | YYCANLKDSRYSGSYYDYW | CQASQDIRFY | LLISDAS | ATYYCQQYDNLPF |
| 7stz | VSGFSLSRYGVH | GMMWGGGNTDY | YYCASSNYVLGYAMDYW | CKSSQSLLNSSNQKNY | LLIYFTS | ADYFCQQHYRTPH |
| 7vux | ASGFAFSSYDMS | ATISGGGRYTYY | YFCASPYGGYFDVW | CRASQSISNF | LLIKYAS | AVYFCQQSNSWPH |

#### 2 On the effect of clustering on docking success rate

As described in the main text, FCC clustering (Rodrigues et al., 2012) is applied at the end of each HADDOCK3 run in order to group together models sharing a consistent fraction of their interfacial contacts. Given the low number of sampled solutions, docking protocols typically give rise to few clusters.

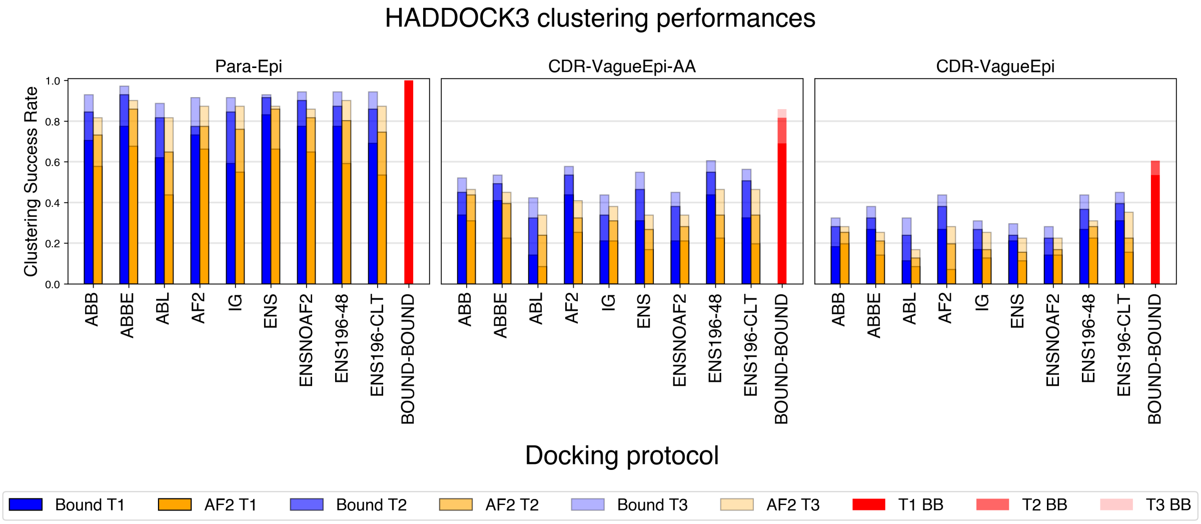


**Supplementary Figure 2:** Cluster-based success rate (CB-SR) of each protocol upon applying FCC Clustering at the end of the workflow. T1, T2, and T3 bars represent the CB-SR over the first one, two, and three clusters, respectively.

Supplementary Fig. 2 shows the cluster-based success rate (CB-SR) over the top 1, top 2 and top 3 clusters for each protocol. Following Ambrosetti et al. (2020), the CB-SR is calculated as the percentage of times in which at least one of the top four models of the top 1, top 2, or top3 clusters is acceptable or more.

According to this definition, CB-SR is very high for the Para-Epi scenario. More specifically, ABBE performs well, reaching CB-SR of $0.78$, $0.93$, and $0.97$ (resp. $0.68$, $0.86$ and $0.90$) over the top 3 clusters when using the bound (resp. AlphaFold2-modelled) antigen. The ABBE protocol outperforms all the other single tool protocols (ABB, ABL, AF2, IG), and even reaches a slightly better accuracy than the best ensemble protocol, ENS, which shows CB-SR values equal to $0.83$, $0.92$, and $0.93$ ($0.66$, $0.86$, and $0.87$ for the AlphaFold2 antigen). There are only two antibodies for which ABBE fails to retrieve an accurate model in the first three clusters, which are 7MSQ and 7F7E. Both structures are not well captured within the ABBE ensemble, with the best model showing a paratope RMSD of $5.29$ and $2.56$ Å. Nevertheless, HADDOCK retrieves two acceptable models for each of these pdb at the rigid-body stage, but these models are not selected by the clustering.

### 3 An analysis of medium and high-quality docking poses

The main text focuses on acceptable quality models according to CAPRI criteria, namely structures that contain at least $10$ % of the native contacts ($F_{nat} > 0.1$) and possess a value of interface-RMSD (I-RMSD) lower than $4.0$ Å or a ligand-RMSD (L-RMSD) below $10$ Å.

Supplementary Fig. 3 shows how the success rate when we consider models of at least medium quality ($F_{nat} > 0.3$ and either I-RMSD $< 2.0$ Å or L-RMSD $< 5.0$ Å). As expected, the success rate is lower than before, but still HADDOCK3 consistently reaches a T1-med SR values around $0.40$ (resp. $0.20$) for the bound (resp. AlphaFold2-modelled) antigen in the Para-Epi scenario. In this scenario, the refinement has an important impact on T1-med SR (resp. T10-med SR) for all protocols, with an average improvement of $0.13$ (resp. $0.15$) when using the bound antigen and of $0.07$ (resp. $0.20$) for runs targeting an AlphaFold2-generated antigen structure.

In the low-information scenario (CDR-VagueEpi-AA) the number of medium or better quality models is lower, with the best protocols (ABBE and all the ensemble protocols) reaching T10-med SR around $0.40$ and $0.23$. Here flexible refinement and energy minimisation are of limited help in re-scoring and improving the success rate, displaying a small improvement only considering the T10 SR of runs targeting the AlphaFold2 antigen ($0.03$).

Supplementary Fig. 4 shows the values of T1-high, T10-high, and T48-high SR, that is, the percentage of times HADDOCK3 generates a high-quality model ($F_{nat} > 0.5$ and either I-RMSD $< 1.0$ Å or L-RMSD $< 1.0$ Å) in the top 1, 10, and 48 structures, respectively. High-quality models are hard to reach with the limited sampling present in our protocols (see main text).

In the good knowledge scenario, protocols obtain a good fraction of high-quality models only when employing the bound structure of the antigen. The best protocols in terms of T10-high SR is ENS with $0.17$. In the CDR-EpiVag-AA scenario (Supplementary Fig. 4b)) most protocols cannot identify more than a handful high-quality poses, with ENS-196-48 and ENS-196-CLT emerging as the best protocols in terms of T10-high SR ($0.17$).

As in the remaining manuscript, also in this case the flexible refinement is very helpful if guided by a good knowledge of the interface (Para-Epi scenario, Supplementary Fig. 4c)), while being detrimental if the restraints are less precise (CDR-VagueEpi-AA scenario, Supplementary Fig. 4d)).

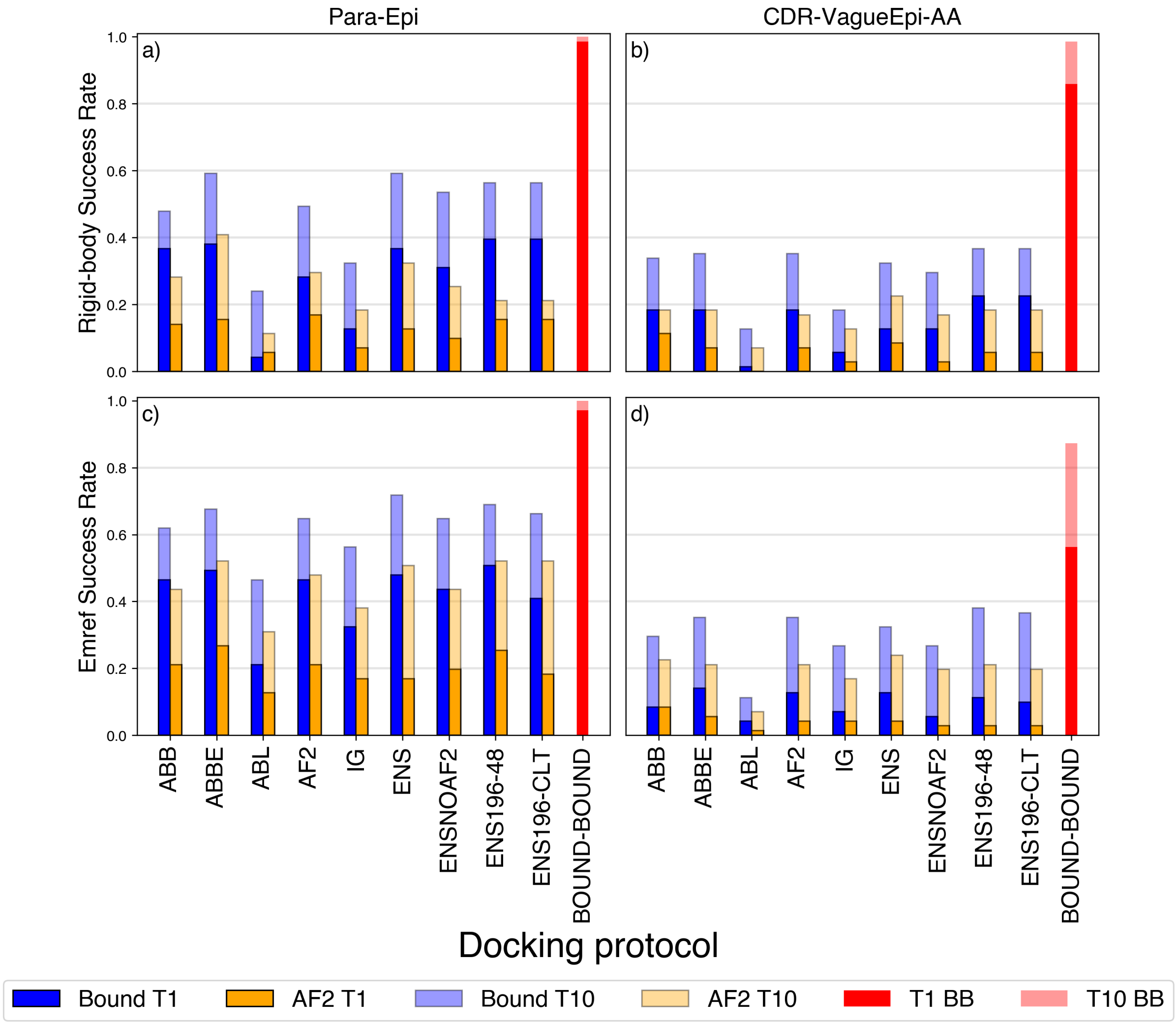


**Supplementary Figure 3:** Success rates for medium quality poses according to CAPRI criteria.

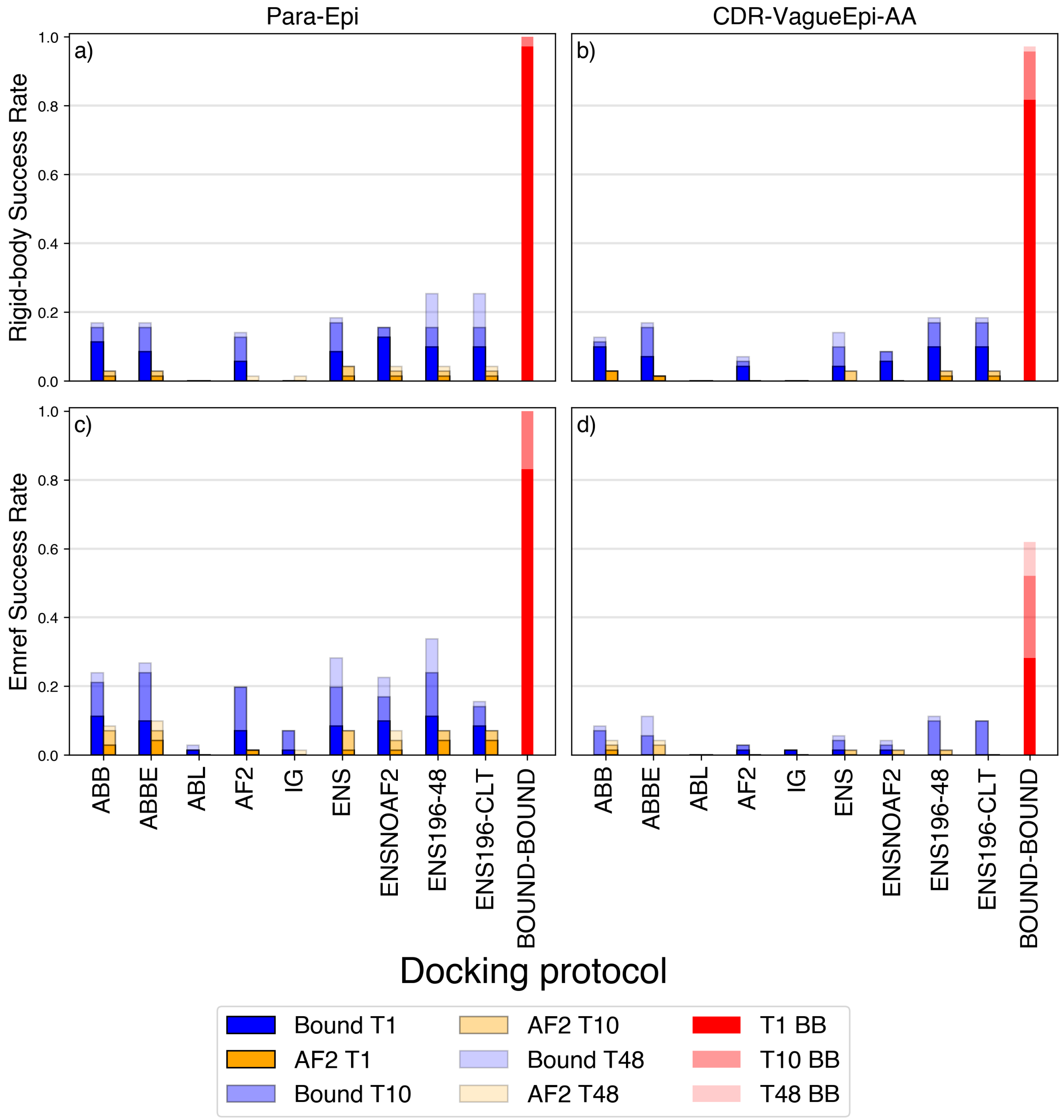


**Supplementary Figure 4:** Success rates for high-quality poses according to CAPRI criteria.

#### 4 Influence of epitope and paratope RMSD on docking quality

In this section we continue the discussion on the importance of epitope and paratope RMSD on docking quality that was begun in the main text. There it was shown that there was a strong correlation between the maximum DockQ score for the top 10 poses and the interface RMSD of the modelled and true antibodies and antigens. We now discuss whether the epitope or paratope RMSD influence docking quality when considered separately.

In Supplementary Fig. 5 and 6, we plot the maximum DockQ score found for the top 10 ranked poses (DockQ-T10) against either the paratope or epitope RMSD after rigid-body and refinement stages, respectively. We further distinguish between different antibody prediction tool. We show both the CDR-Vague-Epi-AA and Para-Epi scenarios, both times using the AlphaFold2 modelled antigen. When comparing the Pearson correlation coefficient $\rho$, in the CDR-VagueEpi-AA scenario it can be seen that the paratope RMSD is a stronger predictor of the maximum DockQ score than the epitope RMSD across all antibody prediction models. The epitope RMSD has a weak anti-correlation with respect to the maximum DockQ score.

We next evaluate the Para-Epi scenario. In this scenario the epitope and paratope RMSD have comparable correlation to the DockQ-T10 score for ABodyBuilder2, but the remaining models have a substantially stronger anti-correlation.

To further evaluate the impact of the epitope RMSD on the docking success rate, we now divide our $71$ antigen structures in $3$ categories, based on the corresponding value of the AlphaFold2 antigen epitope RMSD:

- $61$ "high-quality" structures have a an epitope showing an RMSD lower than $2.0$ Å;
- $9$ "good" antigens possess values of RMSD between $2.0$ and $5.0$ Å;
- $1$ "bad" antigen (7LFB) has an incorrectly modelled epitope (RMSD $>5.0$ Å) showing an epitope RMSD of $5.95$ Å.

We note that HADDOCK3 is never able to generate a good docking pose using the AlphaFold2-modelled antigen structure for 7LFB. This is understandable, as the epitope has been inaccurately predicted by Alphafold2. Supplementary Fig. 7(a-b) shows the average HADDOCK3 SR for top 1 and top 10 models for each protocol over the datasets of good and high quality epitopes. Performances over the $61$ high-quality epitopes are typically much better than those on the $9$ good quality epitopes, especially for the top 1 structures. For example, in Supplementary Fig. 7a) we can observe how the TOP1 SR of ABBE, ENS, ENS196-48 and ENS-196-CLT approaches $60\%$ over the high-quality epitopes.

The trend is still present but slightly less consistent after flexible refinement and energy minimization (Supplementary Fig. 7(c-d)): once the flexibility is introduced performances seem to depend less heavily on the epitope quality for protocols such as ABL and AF2, while a big gap in accuracy remains for the other protocols. As an example, when focusing on high-quality epitopes (Supplementary

Fig. 7(c)) ABBE achieves an outstanding T1-acc SR of $0.64$ ($39$ out of $61$ structures) in the Para-Epi scenario.

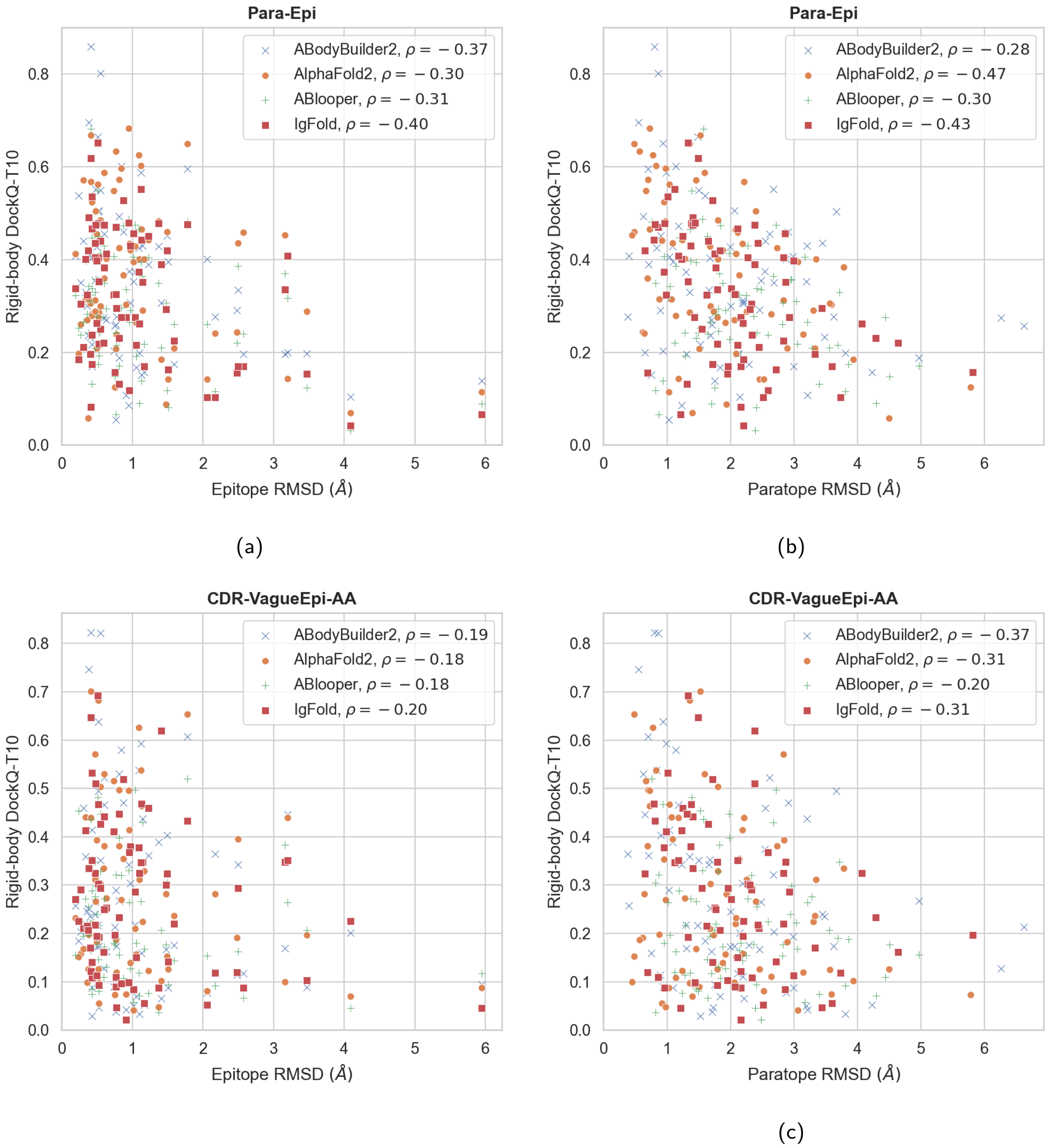


**Supplementary Figure 5:** Panels (a) and (c) are scatter plots showing the maximum DockQ score in the top 10 ranked poses against the epitope RMSD. Panels (b) and (d) show the maximum DockQ score against paratope RMSD. The models are here taken after the rigid-body stage. Points with different colors and shapes correspond to different antibody prediction tools. All plots use the AlphaFold2 modelled antigen. The Pearson correlation coefficient $\rho$ obtained between the two measurements in each plot is given in the legend, broken down by prediction tool.

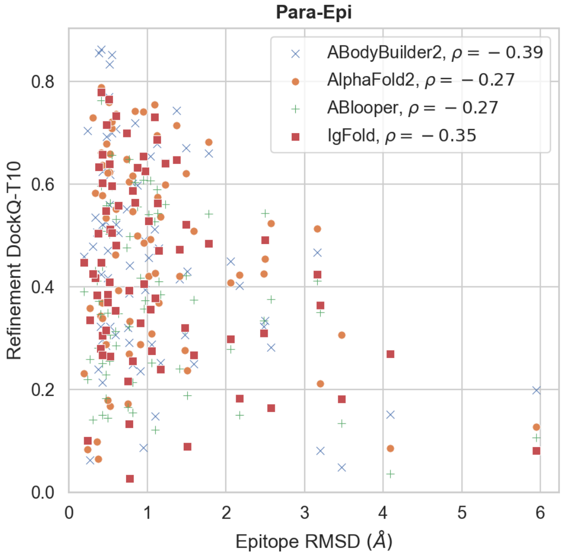


(a)

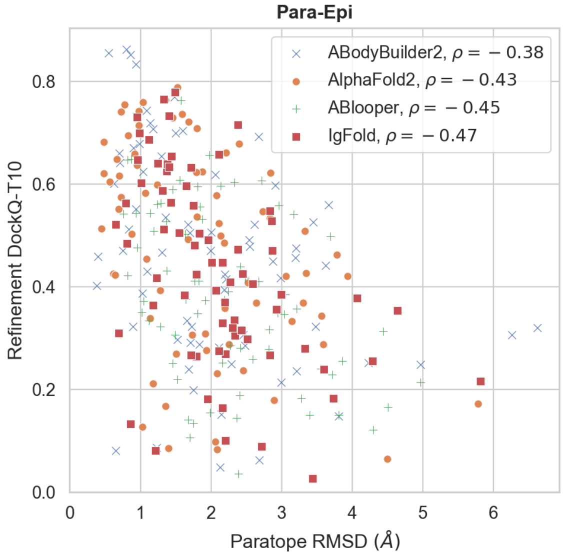


(b)

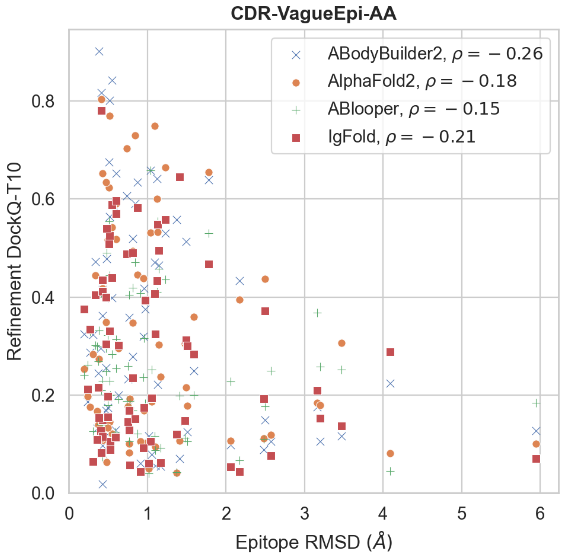


(c)

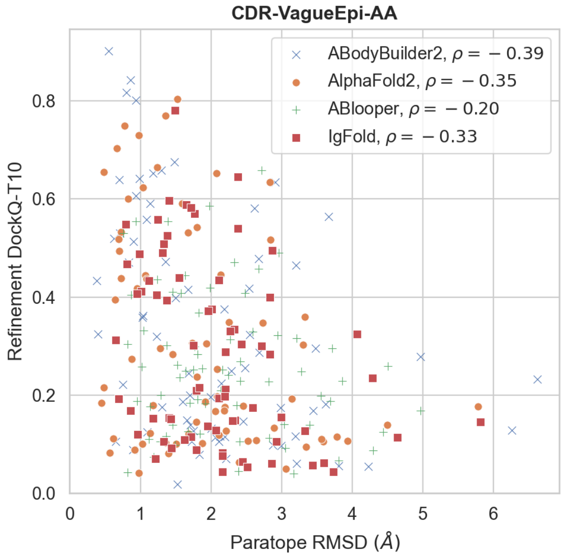


(d)

**Supplementary Figure 6:** Panels (a) and (c) are scatter plots showing the maximum DockQ score in the top 10 ranked poses against the epitope RMSD. Panels (b) and (d) show the maximum DockQ score against paratope RMSD. The models are here taken after the refinement stage. Points with different colors and shapes correspond to different antibody prediction tools. All plots use the AlphaFold2 modelled antigen. The Pearson correlation coefficient $\rho$ obtained between the two measurements in each plot is given in the legend, broken down by prediction tool.

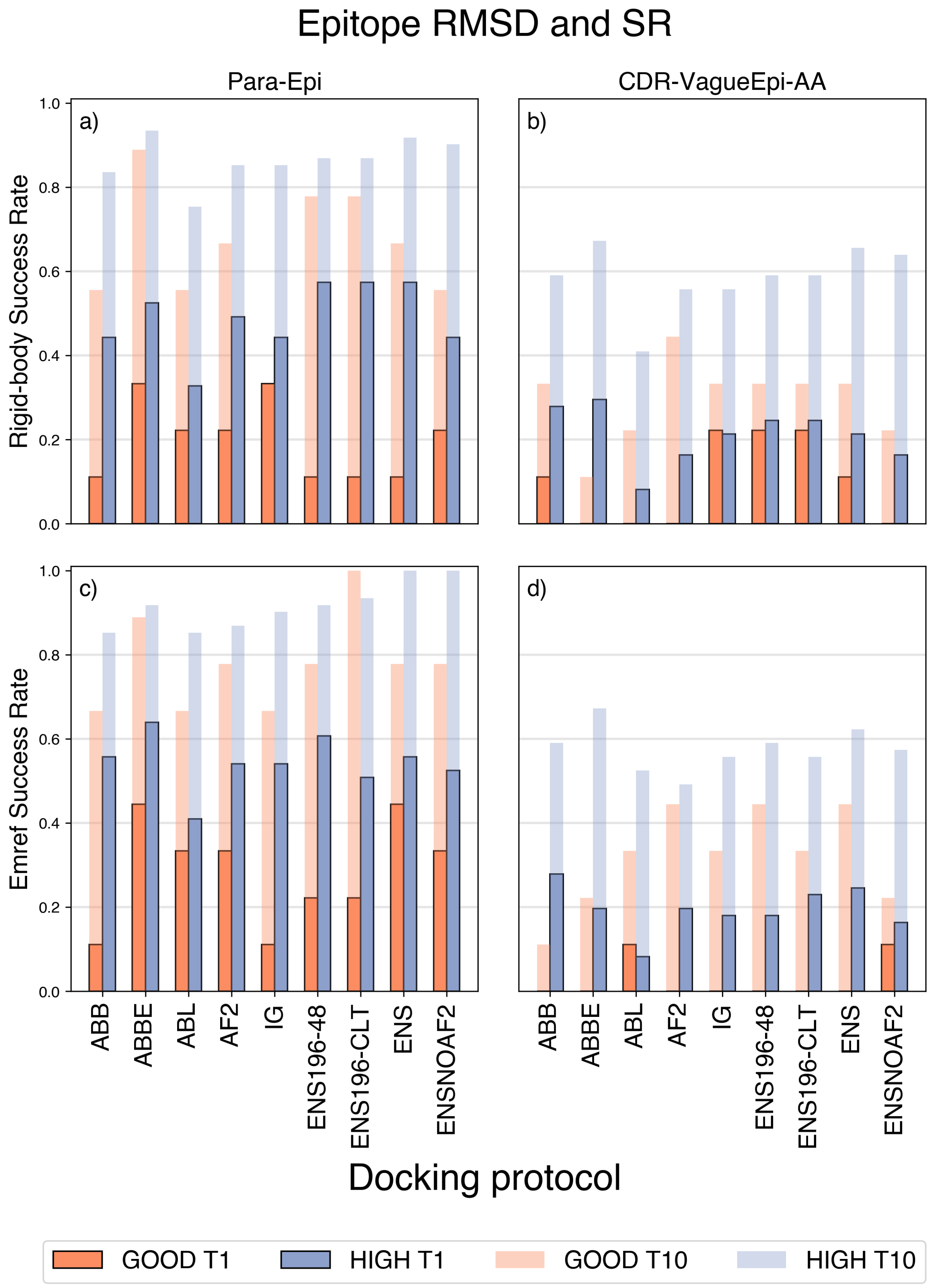


**Supplementary Figure 7:** Relationship between the epitope quality of the AlphaFold2-modelled antigen and docking success rate for the rigid-body and emref stages of the HADDOCK workflow. GOOD T1 and GOOD T10 bars refer to the success rates over the 9 "good" epitopes ( $2.0$ Å $<$ RMSD $< 5.0$ Å) the top 1 and top 10 models, respectively. Analogously, HIGH T1 and HIGH T10 bars refer to the docking success rates for the 61 AlphaFold2-antigens with high quality epitopes (RMSD $< 2.0$ Å). SR values are consistently higher for those runs dealing with a high-quality epitope.

### 5 Epitope and paratope confidence scores as predictors of docking quality

While the previous section showed that the paratope RMSD and epitope RMSD are correlated with the DockQ-T10 score, these measures are only accessible when the true bound structure is known. If this is the case, then there is no reason to use antibody/antigen modelling tools. Some modelling tools have an inbuilt confidence estimator, the pLDDT score for AlphaFold2 (Jumper et al., 2021), and the RMSPE for ABodyBuilder2 (Abanades et al., 2022). In this section we will investigate if these can be used as a predictor of docking quality. If so, these confidence measures could be used to filter out candidate antibodies during an *in silico* virtual screening campaign in order to avoid low quality docked poses.

To begin, we show the correlation between the RMSD of the AlphaFold2 models compared to the true structures and the mean pLDDT, evaluated both on the epitope and on the paratope residues. This is shown in the scatter plots of Supplementary Fig. 8. We find that the mean pLDDT evaluated on the epitope and paratope residues (as defined by the Para-Epi scenario) is strongly anti-correlated with the RMSD on the corresponding region. In Supplementary Fig. 9, we show how removing structures with a mean epitope (paratope) pLDDT lower than a cutoff value modifies the mean epitope (paratope) RMSD over the remaining data set. We also show the mean of the epitope (paratope) RMSD for the removed structures. We can clearly see that the AlphaFold2 pLDDT can be used to remove antibody and antigen structures that are poorly modelled.

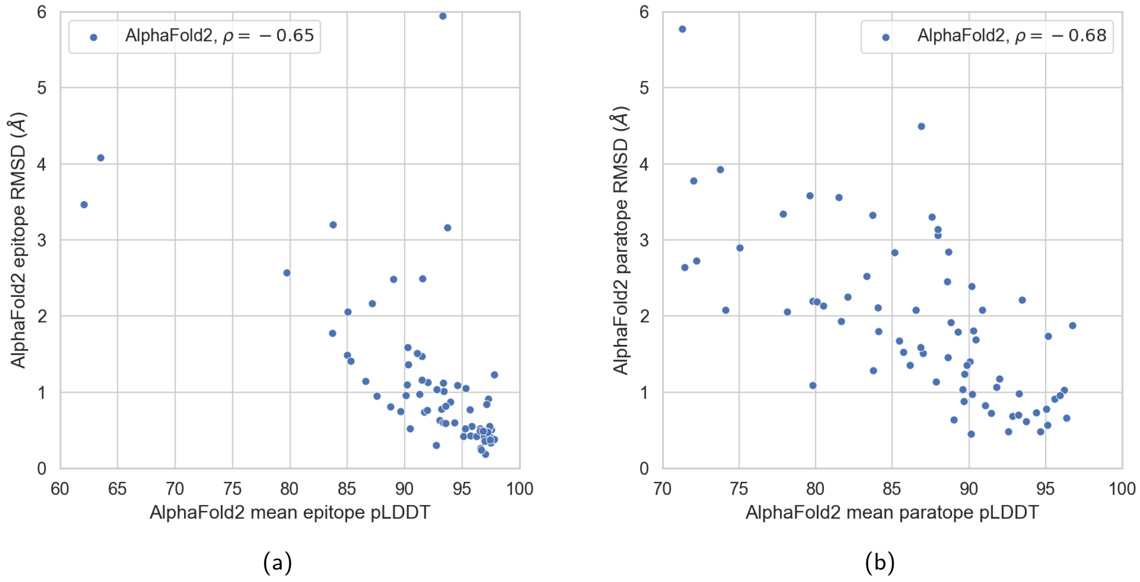


**Supplementary Figure 8:** Scatter plots showing the correlation between the RMSD and mean pLDDT for AlphaFold2 predicted models for the epitope (left) and paratope (right) residues as defined in the Para-Epi scenario. The epitope RMSD and mean pLDDT have a strong anti-correlation ($\rho = -0.65$) as does the paratope RMSD and mean pLDDT ($\rho = -0.68$), showing that the pLDDT can be used to assess model quality for both regions.

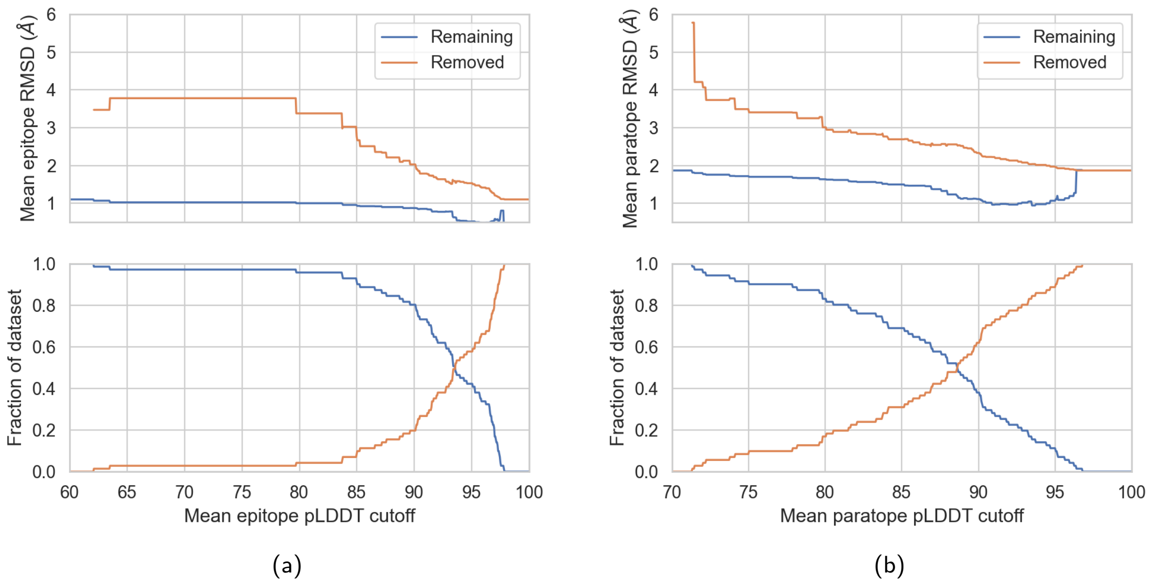


**Supplementary Figure 9:** Plots showing the effect of filtering out structures with small mean pLDDT values on the mean dataset RMSD. The top plot in each panel shows the mean RMSD for both the removed and remaining structures after filtering out structures with a mean pLDDT smaller than a given cutoff. In the left panel we evaluate the mean pLDDT and RMSD over the epitope residues, and in the right panel we evaluate over the paratope residues. In the bottom plot of both panels we show the fraction of the data set has been removed and how much remains.

Next we evaluate the correlation between the mean ABodyBuilder2 RMSPE for the paratope residues and the paratope RMSD between the true antibody and the ABodyBuilder2 model. We plot this in Supplementary Fig. 10. We find a strong correlation of $\rho = 0.72$, indicating that we can also use the mean RMSPE on the paratope to exclude poorly modelled structures. We confirm this in Supplementary Fig. 11 where we show how removing structures with a mean paratope RMSPE lower than a given cutoff changes the mean paratope RMSD of the remaining data set.

Having established that mean epitope pLDDT and mean paratope pLDDT and RMSPE are correlated with the RMSD across the corresponding regions, we investigate the extent to which they are also related to docking success. We will use the DockQ-T10 values introduced in the previous section.

We first show the correlation between DockQ-T10 and AlphaFold2 mean epitope pLDDT in Supplementary Fig. 12. In the CDR-VagueEpi-AA scenario we present the result after rigid-body docking, and the result after flexible refinement and energy minimisation in the Para-Epi scenario. Notably here we use the CDR-VagueEpi-AA definition of epitope residues to compute the mean pLDDT in that scenario, to simulate the information available in that scenario. We break the correlation coefficients down by the antibody model prediction tool. The mean epitope pLDDT is at best weakly correlated with DockQ-T10 in the CDR-VagueEpi-AA scenario, but that the correlation improves in the Para-Epi scenario. This is particularly true for ABodyBuilder2, which obtains a correlation of $\rho = 0.40$. In Supplementary Fig. 13 we illustrate that for ABodyBuilder2 in the Para-Epi scenario filtering out structures with epitopes that have a mean pLDDT lower than a cutoff can remove docking runs with low DockQ-T10, while keeping runs with high DockQ-T10.

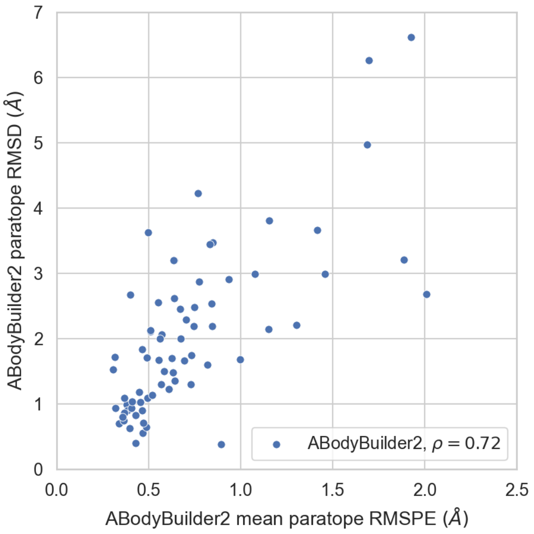


**Supplementary Figure 10:** Plot showing the relationship between the ABodyBuilder2 mean RMSPE score evaluated on the paratope as defined in the Para-Epi scenario and the RMSD between the ABodyBuilder2 modelled paratope and the true structure. We find a strong correlation of $\rho = 0.72$ between the two variables.

The same analysis can be performed on the mean pLDDT for AlphaFold2 modelled paratopes. In Supplementary Fig. 14 we show how DockQ-T10 is correlated with the AlphaFold2 mean paratope pLDDT after the rigid-body stage for the CDR-VagueEpi-AA scenario, and the flexible refinement and energy minimisation stage in the Para-Epi stage. Again we calculate the mean paratope pLDDT by using the antibody residues specified as the paratope to HADDOCK3 in each scenario. There is a weak correlation ($\rho = 0.15$) between the two measurements in the CDR-VagueEpi-AA scenario, and a moderate ($\rho = 0.42$) in the Para-Epi scenario. In Supplementary Fig. 15 we show that the mean paratope pLDDT can be used to remove antibodies with low DockQ-T10 values after docking in the Para-Epi scenario.

To complete this analysis, we evaluate the relationship between the ABodyBuilder2 mean paratope RMSPE and DockQ-T10. The correlation between these two measurements is shown in Supplementary Fig. 16, where we see that in both the CDR-VagueEpi-AA and Para-Epi scenario we obtain moderate anti-correlation with $\rho = -0.35$ and $\rho = -0.45$ respectively. This is sufficient to filter out antibodies which will obtain poor DockQ-T10 values upon docking with HADDOCK3 in both scenarios, as can be seen in Supplementary Fig. 17.

To summarise, both AlphaFold2 paratope pLDDT as well as ABodyBuilder2 paratope RMSPE show a moderate correlation with docking success and could therefore be used as a pre-screening step in docking algorithms using ML-generated antibody and antigen models. The correlation is stronger in the high information Para-Epi scenario, where more accurate definitions of the paratope and epitope residues are available. For docking runs with ABodyBuilder2 in the Para-Epi scenario, the AlphaFold2 epitope pLDDT can also be used to filter out runs which have low docking success.

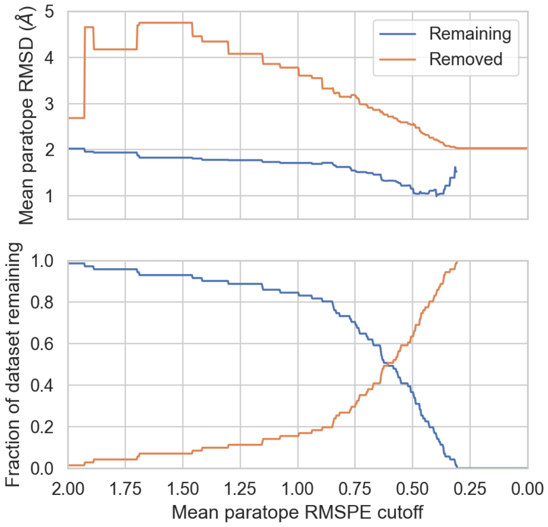


**Supplementary Figure 11:** Plots showing the effect of filtering out antibodies with large paratope RMSPE on paratope RMSD. The top plot shows the mean paratope RMSD for both the removed and remaining structures after filtering out antibodies with a mean paratope RMSPE larger than the given cutoff. The bottom plot shows the fraction of the data set has been removed and how much remains.

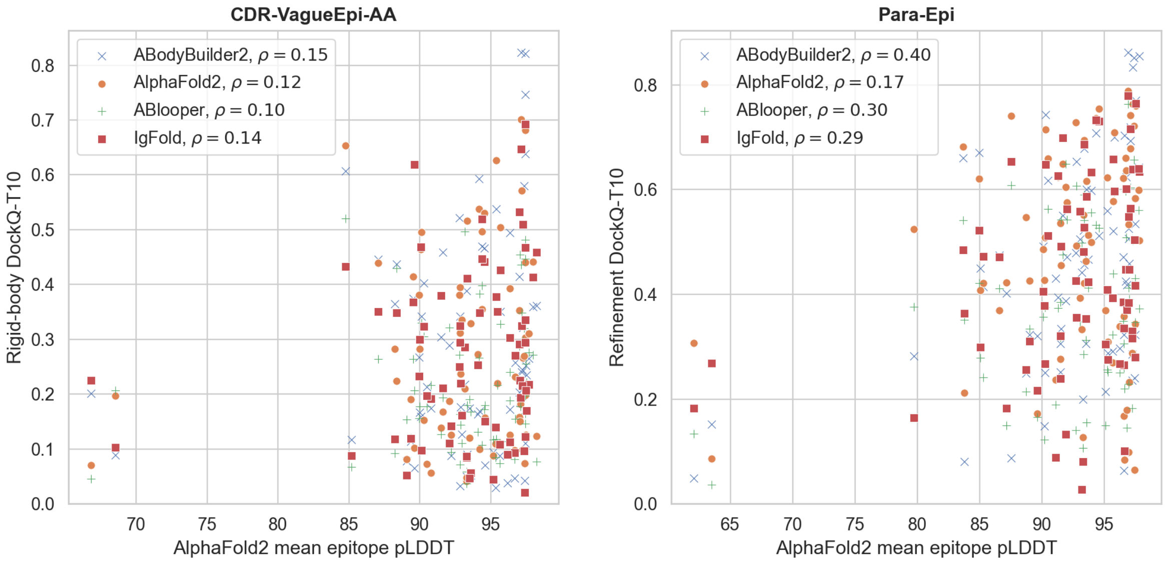


(a) (b)

**Supplementary Figure 12:** Plots showing the maximum DockQ score in the top 10 ranked poses against the AlphaFold2 epitope pLDDT. Panel (a) corresponds to the CDR-VagueEpi-AA scenario, and (b) to the Para-Epi scenario. The CDR-VagueEpi-AA scenario shows results from the rigid-body stage, whereas Para-Epi shows results after the flexible refinement and energy minimisation stage. To compute the mean epitope pLDDT, the epitope residues are taken to be those as specified to HADDOCK3 in the given scenario. Points with different colors and shapes correspond to different antibody prediction tools. The Pearson correlation coefficient $\rho$ obtained between the two measurements in each plot is given in the legend, broken down by prediction tool.

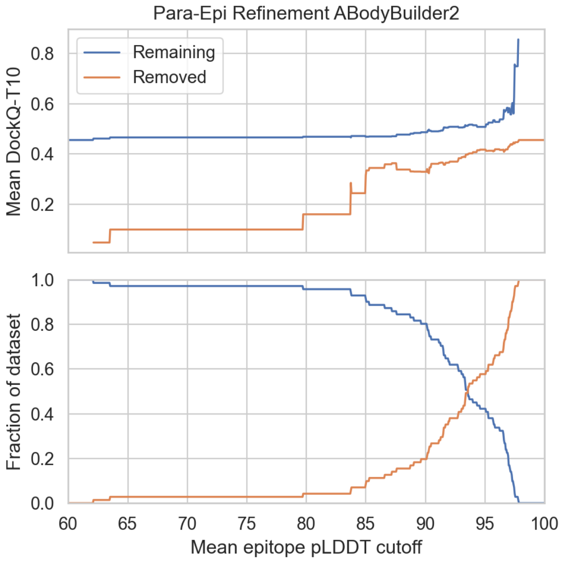


**Supplementary Figure 13:** Plots showing the effect on DockQ-T10 after filtering out complexes based on epitope pLDDT in the Para-Epi scenario, after flexible refinement and energy minimisation and using the ABodyBuilder2 modelled antibody. The top plot shows the mean DockQ-T10 for both the removed and remaining complexes after filtering out complexes with an antigen with mean epitope pLDDT smaller than the given cutoff. The bottom plot shows the fraction of the data set has been removed and how much remains.

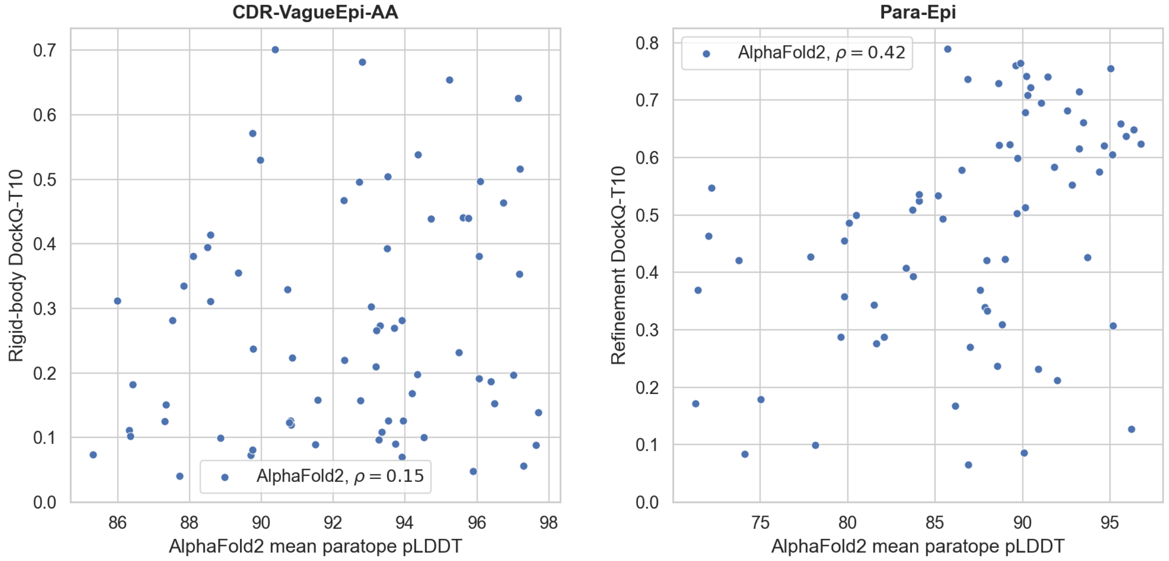


(a) (b)

**Supplementary Figure 14:** Plots showing the maximum DockQ score in the top 10 ranked poses against the AlphaFold2 paratope pLDDT. Panel (a) corresponds to the CDR-VagueEpi-AA scenario, and (b) to the Para-Epi scenario. The CDR-VagueEpi-AA scenario shows results from the rigid-body stage, whereas Para-Epi shows results after the flexible refinement and energy minimisation stage. To compute the mean paratope pLDDT, the paratope residues are taken to be those as specified to HADDOCK3 in the given scenario. Both plots use the AlphaFold2 modelled antigen. The Pearson correlation coefficient $\rho$ obtained between the two measurements in each plot is given in the legend.

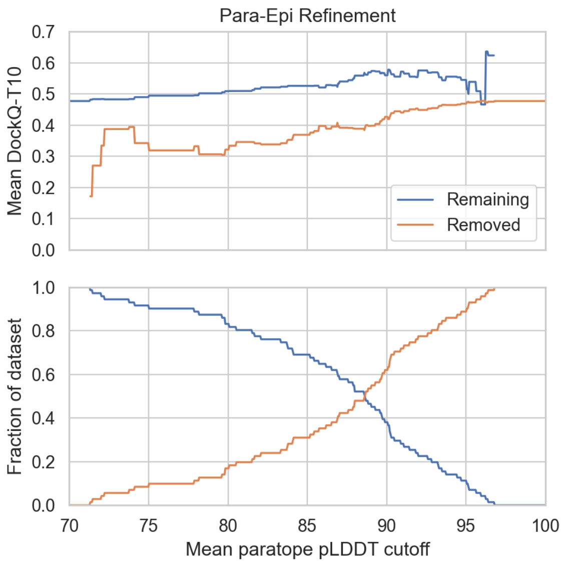


**Supplementary Figure 15:** Plots showing the effect on DockQ-T10 after filtering out complexes based on paratope pLDDT in the Para-Epi scenario, after flexible refinement and energy minimisation and using the AlphaFold2 modelled antibody. The top plot shows the mean DockQ-T10 for both the removed and remaining complexes after filtering out complexes with an antibody with mean paratope pLDDT smaller than the given cutoff. The bottom plot shows the fraction of the data set has been removed and how much remains.

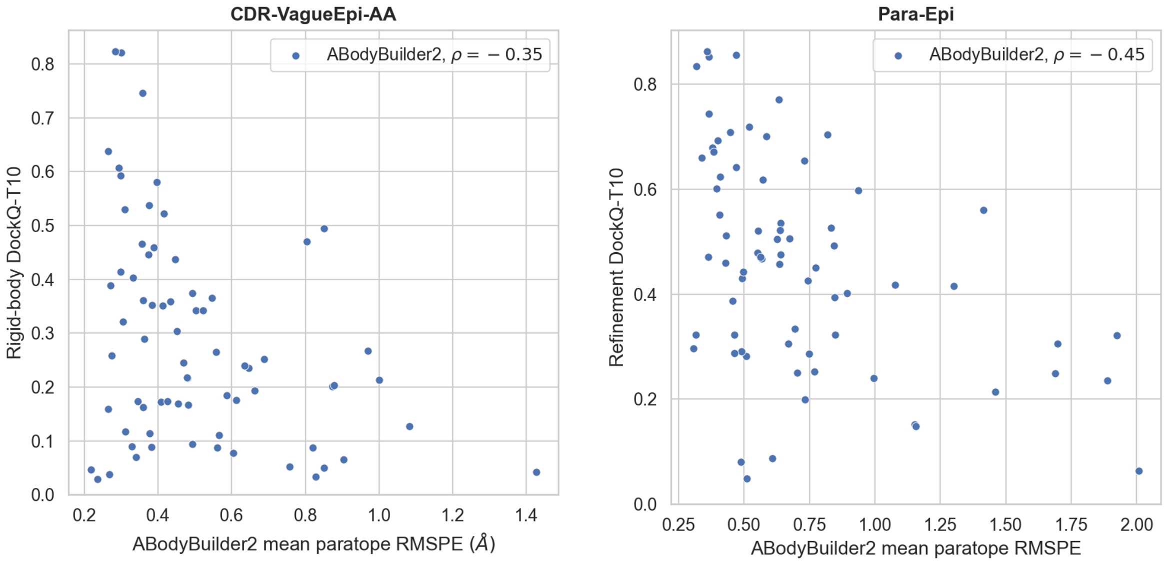


(a) (b)

**Supplementary Figure 16:** Plots showing DockQ-T10against the ABodyBuilder2 mean paratope RMSPE. Panel (a) corresponds to the CDR-VagueEpi-AA scenario, and (b) to the Para-Epi scenario. The CDR-VagueEpi-AA scenario shows results from the rigid-body stage, whereas Para-Epi shows results after the flexible refinement and energy minimisation stage. To compute the mean paratope RMSPE, the paratope residues are taken to be those as specified to HADDOCK3 in the given scenario. Both plots use the AlphaFold2 modelled antigen. The Pearson correlation coefficient $\rho$ obtained between the two measurements in each plot is given in the legend.

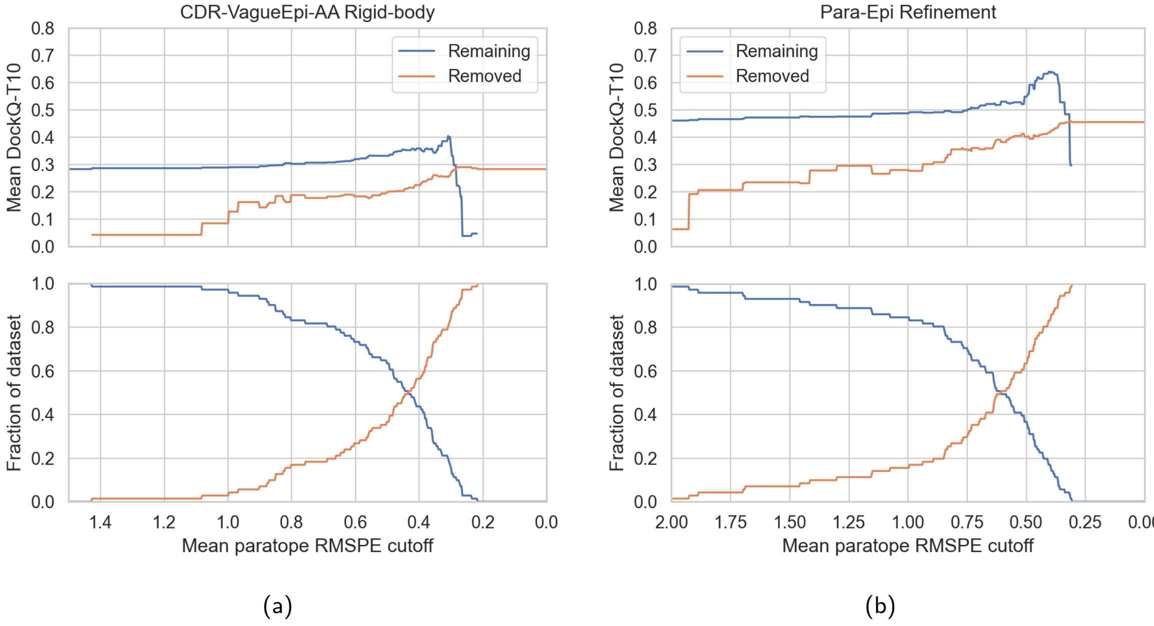


**Supplementary Figure 17:** Plots showing the effect on DockQ-T10 after filtering out complexes based on mean paratope RMSPE. In both panels the top plot shows the mean DockQ-T10 for both the removed and remaining complexes after filtering out complexes with an antibody with mean paratope RMSPE larger than the given cutoff. The bottom plot shows the fraction of the data set that has been removed and how much remains. The left panel shows the DockQ-T10 results after the rigid-body stage in the CDR-VagueEpi-AA scenario, whereas the right panel shows the results after the flexible refinement and energy minimisation stage after the Para-Epi scenario. All plots use the AlphaFold2 modelled antigen. The residues used to calculate the mean paratope RMSPE in all plots are those specified as the paratope in the given scenario to HADDOCK3.

### 6 Energy minimisation of rigid-body solutions increases the top 1 success rate

In the main text we discuss the importance of interface refinement for the analysed protocols, highlighting how this CPU-intensive stage can be avoided, especially in the CDR-VagueEpi-AA scenario.

We then applied a short, unexpensive (see Table 2), HADDOCK energy minimisation step to the rigid-body docking models. The results suggest that this energy minimisation is beneficial for the T1-acc SR (see Supplementary Fig. 18), while it seems to have a slightly negative effect on the T10-acc SR, in particular in the CDR-VagueEpi-AA scenario. ABB, ABBE, ENS196-48, ENS196-CLT protocols show substantial (higher than $0.05$) improvements in success rates in the Para-Epi scenario, notably reaching a value of T1-acc close to $0.75$ without any explicit refinement.

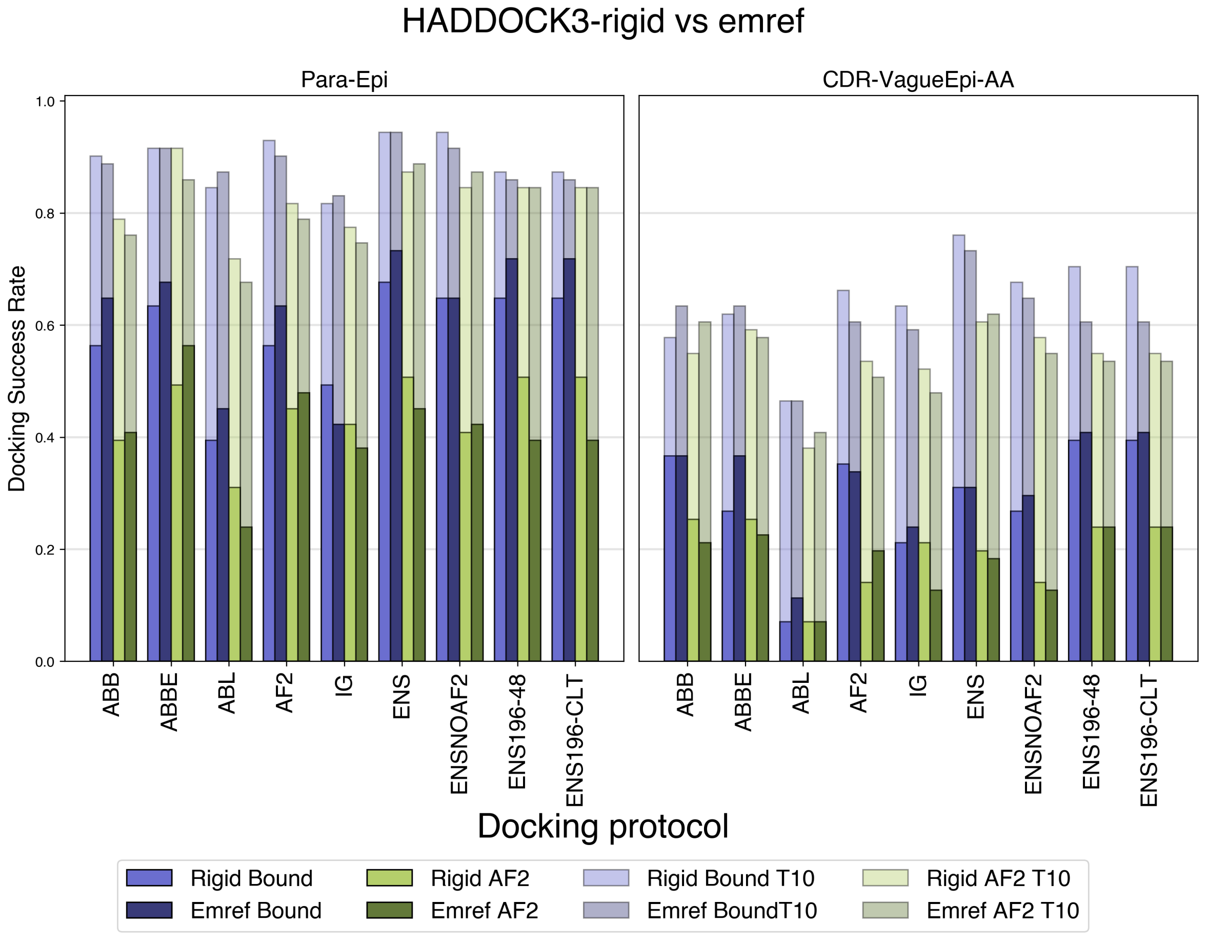


**Supplementary Figure 18:** Comparison of success rates between rigid-body docking models (RIGID) and energy-minimised rigid-body models (EMREF).

### 7 Protocol timings

Alongside accuracy, for high-throughput docking applications execution speed is also an important performance metric of docking algorithms. Table 2 shows the average execution time (in CPU hours) for the rigid-body and flexible refinement stages of the HADDOCK3 protocol, alongside the average HADDOCK3 and ZDOCK execution time. The pure rigid-body, FFT-based ZDOCK is faster than HADDOCK3 when considering total execution time. Considering only rigid-body energy minimisation in the Para-Epi scenario however, the two software show similar execution speed. In the main text, we show that HADDOCK3 already produces accurate predictions at this stage, potentially eliminating the need for the resource intensive flexible refinement. It is important to note the impact of the restraints on HADDOCK3 execution time: when accurate information is available (Para-Epi scenario), few restraints are imposed and the minimisation is fast. CDR-VagueEpi-AA runs on the other hand are slower than CDR-VagueEpi runs, as defining two sets of residues as active implies imposing a higher number of restraints.

**Supplementary Table 2:** Average execution time (in CPU hours) for Default docking protocols (excluding ENS-196-48 and ENS-196-CLT).

| **Software** | Para-Epi | CDR-VagueEpi | CDR-VagueEpi-AA |
|---|---|---|---|
| **ZDOCK** | 0.24 | 0.25 | 0.25 |
| **HADDOCK rigid** | 0.33 | 0.63 | 1.09 |
| **HADDOCK flexref** | 2.47 | 3.98 | 6.04 |
| **HADDOCK emref** | 0.20 | 0.30 | 0.43 |
| **HADDOCK full** | 3.12 | 5.02 | 7.67 |

John Jumper, Richard Evans, Alexander Pritzel, Tim Green, Michael Figurnov, Olaf Ronneberger, Kathryn Tunyasuvunakool, Russ Bates, Augustin Žídek, Anna Potapenko, Alex Bridgland, Clemens Meyer, Simon A. A. Kohl, Andrew J. Ballard, Andrew Cowie, Bernardino Romera-Paredes, Stanislav Nikolov, Rishub Jain, Jonas Adler, Trevor Back, Stig Petersen, David Reiman, Ellen Clancy, Michal Zielinski, Martin Steinegger, Michalina Pacholska, Tamas Berghammer, Sebastian Bodenstein, David Silver, Oriol Vinyals, Andrew W. Senior, Koray Kavukcuoglu, Pushmeet Kohli, and Demis Hassabis. Highly accurate protein structure prediction with AlphaFold. *Nature*, 596(7873):583–589, 8 2021. ISSN 0028-0836. doi: 10.1038/s41586-021-03819-2. URL `https://www.nature.com/articles/s41586-021-03819-2`.

João PGLM Rodrigues, Mikaël Trellet, Christophe Schmitz, Panagiotis Kastritis, Ezgi Karaca, Adrien SJ Melquiond, and Alexandre MJJ Bonvin. Clustering biomolecular complexes by residue contacts similarity. *Proteins: Structure, Function, and Bioinformatics*, 80(7):1810–1817, 2012.
